# Comparison of cell response to chromatin and DNA damage

**DOI:** 10.1101/2023.01.17.524424

**Authors:** Artyom Luzhin, Priyanka Rajan, Alfiya Safina, Katerina Leonova, Aimee Stablewski, Jianmin Wang, Mahadeb Pal, Omar Kantidze, Katerina Gurova

## Abstract

DNA-targeting drugs may damage DNA or chromatin. Many anti-cancer drugs damage both, making it difficult to understand their mechanisms of action. Using molecules causing DNA breaks without altering nucleosome structure (bleomycin) or destabilizing nucleosomes without damaging DNA (curaxin), we investigated the consequences of DNA or chromatin damage in normal and tumor cells. As expected, DNA damage caused p53-dependent growth arrest followed by senescence. Chromatin damage caused higher p53 accumulation than DNA damage; however, growth arrest was p53-independent and did not result in senescence. Chromatin damage activated the transcription of multiple genes, including classical p53 targets, in a p53-independent manner. Although these genes were not highly expressed in basal conditions, they had chromatin organization around the transcription start sites (TSS) characteristic of most highly expressed genes and the highest level of paused RNA polymerase. We hypothesized that nucleosomes around the TSS of these genes were the most sensitive to chromatin damage. Therefore, nucleosome loss upon curaxin treatment would enable transcription without the assistance of sequence-specific transcription factors. We confirmed this hypothesis by showing greater nucleosome loss around the TSS of these genes upon curaxin treatment and activation of a p53-specific reporter in p53-null cells by chromatin-damaging agents but not DNA-damaging agents.

## Introduction

DNA-binding small molecules have been used in medicine and biology for a long time. They showed multiple clinical and biological effects, including anti-cancer, anti-infective, and anti-inflammatory activities. Most of their effects are attributed to their ability to cause DNA damage (DD), i.e., alter DNA chemical structure. The effects of many of these molecules on the structure and function of chromatin in cells and cell-free conditions were observed long ago (1), (2); however, chromatin-related effects were not investigated in depth because their biological significance and impact on their anti-cancer activity were unknown. Recently, the chromatin-related effects of DNA binding drugs have been getting more attention, with a focus on separating the cellular consequences of DNA damage from the chromatin alterations caused by these molecules. Two pivotal studies published in 2013 demonstrated that anti-cancer agents from anthracycline group cause histone eviction from chromatin and that this effect does not correlate with DNA damaging activity of these molecules (3,4). It was proposed that histone eviction may result from their effects on DNA topology (4) and may have implications for the mechanisms underlying cell killing during cancer chemotherapy (3,4).

Our group discovered (5) the curaxins, candidate anti-cancer agents, and defined their mechanism of action (6). Curaxins are chemically different from anthracyclines (i.e., carbazole vs. anthraquinone derivatives); however, there are similarities in their DNA binding. Curaxins are the appropriate tools for distinguishing between chromatin-mediated and DNA-damaging effects. These molecules do not damage DNA in mammalian cells but efficiently evict histones from chromatin, an effect we named “chromatin damage” (CD) (7,8). Similar to anthracyclines, curaxins have demonstrated anti-tumor activity in multiple preclinical models (i.e., have higher toxicity to tumor cells than normal cells), suggesting that tumor cells are more sensitive to CD than normal cells (5,9–14). The reason behind this phenomenon is unclear.

Curaxin binding to DNA causes the lengthening of the double helix, reduces DNA flexibility and its negative charge (curaxins have a small positive charge), and competes for minor groove binding with histones. However, whether all these effects or just some are important for CD and what structural features of these small molecules define the strength of their effect on chromatin are currently unknown. We previously characterized the effect of several known anti-cancer agents differing in their ability to bind DNA on DNA, chromatin, and tumor cell viability to understand why some compounds cause CD and others cause DD or both CD and DD (8). We found that only compounds binding DNA directly cause CD, whereas compounds chemically modifying DNA without direct, stable DNA binding (e.g., bleomycin causes DNA breaks through nuclease-like activity (15–17)), or compounds incorporated into DNA as modified nucleotides, or compounds binding and inhibiting DNA enzymes that use DNA as a substrate (e.g., topoisomerase inhibitor etoposide does not bind DNA per se) do not cause direct CD. Moreover, we observed that CD activity of small molecules had a higher impact on cytotoxicity than DD activity (8). In contrast to CD, the cellular stress response to DD is well-studied; however, in most studies molecules used to cause DD were also causing CD, but this effect was not considered. This suggests that some effects attributed to DD may, in fact, be caused by CD.

A major long-term side effect of anthracycline chemotherapy is cardiotoxicity (18). Recently, Neefjes group (19) demonstrated that cardiotoxicity is caused by compounds that induce strong DD (e.g., doxorubicin), whereas similar compounds with very weak DD activity (e.g., aclarubicin) have similar anti-cancer effects without cardiotoxicity, suggesting that although cardiotoxicity is a consequence of DD, its anti-cancer activity is not (19). Because of these data, it is even more important to separate the cellular consequences of DD and CD to be able to design and use drugs with maximal anti-cancer activity and minimal long-term toxicity. Here, we used agents that, in the short-term, predominantly cause DD (i.e., DNA breaks) or CD (i.e., histone eviction from chromatin without DNA breaks) to distinguish between the cellular effects caused by CD and those resulting from DD. As inducer of CD we used curaxins CBL0137. As DD inducers we selected glycopeptide antibiotics from Streptomyces verticillus belonging to the bleomycin family. These compounds cause double-strand (ds) DNA breaks (rev. in (20)). The crystal structure of bleomycin with DNA shows that it binds DNA and acts like a nuclease. It has a metal binding domain and causes site-selective cleavage of duplex DNA at 5’-GT/C sites after oxygenation (15–17). Importantly, bleomycin does not digest nucleosomal DNA but attacks only nucleosome-free DNA (21). Therefore, it does not cause CD.

## Materials and Methods

### Reagents

CBL0137 was provided by Incuron, Inc and dissolved in dimethyl sulfoxide at 20 mM. Bleomycin solution for injection was provided by Roswell Park Pharmacy (leftover from clinical use) as 30 mg/ml solution. In most experiments we used Zeocin (trade name of phleomycin D1, antibiotic structurally similar to bleomycin) after confirming activity identical to bleomycin. It was purchased from Thermo Fisher Scientific (Grand Island, NY) as 50 mg/ml) solution. Resazurin sodium salt was purchased from Thermo Fisher Scientific. Resazurin was dissolved in Dulbecco’s phosphate-buffered saline (DPBS, pH 7.4) at 0.15 mg/ml (10X solution).

### Cells

The HT1080 lung fibrosarcoma cell line was originally from the American Type Culture Collection. Its authenticity was confirmed using short tandem repeat analysis (100% match). HT1080 cells were maintained in Dulbecco’s Modified Eagle Medium (DMEM) supplemented with 5% (v/v) FBS and 100 units/ml penicillin. Primary human neonatal dermal fibroblasts (NDFs) were obtained from AllCells, LLC (Alameda, CA), as a pool of three separate donors. NDFs were maintained in DMEM supplemented with 10% (v/v) FBS, 100 units/mL penicillin, 100 μg/mL streptomycin, and 2 mM L-glutamine. Cells were cultured in a tissue culture incubator at 37°C and 5% CO_2_.

### Plasmids

The p53-Luc lentiviral plasmid and GSE56 LXSN vector were previously described (22,23). Lentivirus production and transduction were done as described (22). Transfection was performed using Lipofectamine™ 3000 (Invitrogen, Thermo Fisher Scientific (Grand Island, NY)) according to manufacturer protocol.

### Deletion of p53 in cells

Four single-guide RNA (sgRNA) sequences were used to target intron 1, exon 3, exon 4 and intron 9 of human p53 to knock out the gene in HT1080 and NDF cells, as previously described (24,25): ACTTCCTGAAAACAACGTTCTGG (e3), GAGCGCTGCTCAGATAGCGATGG (e4), TCTGCAGGCCCAGGTGACCCagg (i1.1), and GAAACTTTCCA CTTGATAAGagg (i9). Crispr RNA (crRNA) and tracer RNA (trRNA) were purchased from IDT DNA Technologies (Coralville, IA) and resuspended to 160 μM each. crRNA and trRNA (1:1) were complexed using touchdown polymerase chain reaction (PCR) (IDT DNA Technologies), and Cas9 3NLS protein (IDT DNA Technologies) was added to make functional ribonucleoprotein (RNP). RNP was added to 3×10^6^ cells suspended in Opti-MEM medium (ThermoFisher Scientific). The cell mixture was electroporated using a NEPA21 electroporator (Bulldog Bio, Portsmouth, NH) and then transferred into 6-well plates containing complete medium without antibiotics (2 ml). Three electroporation regimens differing in voltage and time of stroke were used with similar results.

After recovery from electroporation, the cells were expanded and frozen. Cells from each electroporation regimen were plated in two plates. One plate was left untreated, and the second plate was treated with 1 μM CBL0137 for 24 hr to induce p53. Non-electroporated cells were used as control cells with wild type p53. The next day, cells were stained for p53, and the number of p53-positive cells was assessed by flow cytometry (Supplementary Figure S3).

In parallel, cells were plated in 96-well plates at 1 cell/well. After colony formation (7-10 days), wells were inspected using microscopy and wells with no colonies or more than one colony were excluded. Cells from other wells were trypsinized and replated into three new 96 well plates. One plate was used as a reference plate and two other plates were either left untreated or treated with 1 μM CBL0137 for 24 hr. These two plates were stained for p53 after that, and the p53-positive and p53-negative clones were identified using fluorescence microscopy. All p53-negative clones were pooled. Loss of p53 was confirmed by western blotting under basal and CBL0137-treated conditions.

### Luciferase reporter assay

Cells were plated in 96-well plates at 5,000 cells per well in 100 μL medium. The next day, 100 μL medium with drugs was added to the cells for 24 hr. Luciferase activity was measured using a BrightGlo kit from Promega (Madison, WI).

### Viability assay

Cells were plated in 96-well plates at 1,000 cells per well in 100 μL medium. The next day, 100 μL medium with drugs was added to the cells for 72 hr. The effects of treatment were determined by adding 20 μL 10× resazurin solution for 4–8 hr. Fluorescence (560/590 nm) was measured using a multi-plate reader. Alternatively, cells were plated in 384-well plates. The next day, the cells were treated with different drug concentrations in triplicate. Cell numbers were counted every 12 hr using the Cytation 5 Cell Imaging Multimode Reader with Gen5 ImagePrime software.

### Induction of senescence

Cells were plated at 5×10^5^ per 10-cm plate and next day treated with CBL0137 or bleomycin for 24 hr. The compounds were washed off after treatment, and the cells were cultured for 10 days with medium changes every 72 hr. After 10 days, cells were trypsinized and plated for colony formation assay, acidic beta-galactosidase staining or measurement of IL-6 and IL-8 in culture medium.

### Colony formation assay

10 days after treatment with CBL0137 or bleomycin, the cells were trypsinized, seeded at 100 cells per 10-cm plate, and allowed to grow for approximately 2 weeks. Cells were washed with PBS, and stained with 1% methylene blue in methanol. The experiment was performed in triplicate for each condition, and colonies of ≥ 50 cells were counted by light microscopy at 10× magnification.

### Senescence-associated β-galactosidase staining

10 days after treatment with CBL0137 or bleomycin, the cells were trypsinized and seeded at 12×10^3^ cells per well in 12-well plates. After 24 hr, the cells were fixed, and stained for acidic β-galactosidase activity as previously described ((26)). Briefly, cells were washed twice in PBS, fixed for 5 min at room temperature with 2% (v/v) paraformaldehyde and 0.2% (v/v) glutaraldehyde, washed with PBS and then incubated with freshly prepared SA-β-Gal staining solution (1mg/ml X-Gal, Thermo Fisher Scientific) dissolved in N,N-dimethylformamide (Sigma-Aldrich), 100 mM citric acid/sodium phosphate buffer (pH 6.0) containing 2 mM MgCl_2_, 150 mM NaCl, 5 mM K_3_Fe(CN)_6_, and 5 mM K_4_Fe(CN)_6_. Staining was performed at 37°C in a non-CO_2_ incubator for up to 18 hr until the X-Gal product was visible. The reaction was stopped by washing the cells with PBS.

### Measurement of IL-6 and IL-8 in culture medium

IL-6 and IL-8 were measured in culture medium using enzyme-linked immunosorbent assays (ELISA). 10 days after treatment with CBL0137 or bleomycin, the cells were trypsinized and seeded at 12×10^3^ cells per well in 12-well plates in 1 ml complete medium. After 72 hr, the conditioned medium was collected and frozen at −80°C. The concentrations of IL-6 and IL-8 were measured in triplicate with the DuoSet ELISA Development System (R&D Systems, Minneapolis, MN) according to the manufacturer’s instructions. Cytokine levels were normalized to the cell number for each condition.

### Comet assay (single-cell gel electrophoresis)

Glass slides were coated with 1% low melting point agarose (LMPA, Type 1A low electroendosmosis agarose (Sigma-Aldrich, St. Louis, MO, cat # A0169) in MilliQ water (Sigma-Aldrich, St. Louis, MO). Once solidified, two wells were created using the top of a 1-ml pipette tip. The slides were left at room temperature overnight in a sealed box to avoid evaporation. Treated and untreated cells were trypsinized and then resuspended in 37°C 1% LMPA at a concentration of 1.1 ×10^6^ cells/ml (50,000 cells per 90 μl per sample). The cell-agarose suspension (90 μL) was added to each well of the slides, which were chilled on ice. After a gel formed, a layer of 0.5% LMPA (100 μL) was added to the wells and allowed to solidify.

The slides were placed in freshly made lysis buffer (2.5 M NaCl, 100 mM EDTA, 10 mM tris base, pH 10, 1% Triton X-100) for 2.5 hr at 4°C, protected from light. The slides were then placed in an alkaline solution (12 g/L NaOH, 500 mM EDTA) at 4°C for 30 min, protected from light. After removing the alkaline solution, the slides were immersed twice in 1× TBE electrophoresis buffer for 5 min each. Electrophoresis was performed at 24 V (0.74 V/cm) for 30 min. After electrophoresis, the slides were washed thrice in prechilled water for 2 min each. The slides were dehydrated with ice-cold 70% ethanol for 5 min, allowed to air dry, and then stained with Vista Green (Cell Biolabs, San Diego, CA, cat # 235005) diluted 1:10,000 in TE buffer. Comets were scored using the OpenComet plugin from ImageJ. The percentage of tail DNA across various samples was determined. The fraction of cells with comets was calculated to determine the prevalence of DNA damage across the samples.

### Immunoblotting

Western blotting was performed as described (27) using the following antibodies

| Antibody | Company | Catalog # | Dilution |
| --- | --- | --- | --- |
| P53 DO-1 | Santa Cruz<br>Biotechnology | sc-126 | 1:1000 (WB),<br>1:200 (IF) |
| P21 F-5 | Santa Cruz<br>Biotechnology | sc-6246 | 1:200 (WB) |
| P21 12D1 | Cell Signaling<br>Technologies | 2947S | 1:2000 (WB) |
| MDM2 SMP14 | Santa Cruz<br>Biotechnology | sc-965 | 1:200 (WB) |
| SSRP1 | BioLegend | 609702 | 1:4000 (WB) |
| SPT16 | BioLegend | 607002 | 1:1000 (WB) |
| γH2AX | Cell Signaling<br>Technologies | 9718S | 1:1000 (WB) |
| Beta-Actin | Sigma-Aldrich | A3854 | 1:30000 (WB) |
| Anti-mouse HRP | Cell Signaling<br>Technologies | sc-516102 | 1:5000 (WB) |
| Anti-rabbit HRP | Santa Cruz<br>Biotechnology | sc-2357 | 1:4000 (WB) |
| Anti-mouse Alexa-Fluor488 | Invitrogen | A21202 | 1:500 (IF) |

### P53 staining for flow cytometry

Cell suspensions (1×10^5^ cells in 300 μL 3% BSA in PBS) or adherent cells on a p60 plate were fixed in 1 ml 4% paraformaldehyde (PFA) for 10 min at room temperature. After fixing, the cells were washed thrice with PBS, blocked with 3% BSA in PBS containing 0.1% Triton X-100 for 1 hr, and stained with primary and secondary antibodies for 1 hr each. The cells were washed thrice with 0.1% Triton X-100 in PBS after each antibody incubation. Resuspended cells were analyzed by flow cytometry, and the adherent cells were assessed using a Zeiss AxioVision A1 fluorescent microscope.

### 5-Ethynyl-2’-deoxyuridine (EdU) incorporation assay

Cells were treated with 1 μM CBL0137 or 500 μg/ml bleomycin for 24 hr and then incubated with 10μM of EdU for 2 hr. After treatment, the cells were fixed with 4% PFA for 15 min. EdU incorporation was detected with the Click-IT Plus EdU Flow Cytometry Assay (Invitrogen, cat#10632) using the manufacturer’s staining protocol.

### 5-Ethynyl uridine (EU) incorporation assay

Cells were treated with CBL0137 (0, 0.25, 0.5, and 1 μM) for 3 hr, with EdU (0.2 mM) added for the last hour. After treatment, the cells were fixed with 4% PFA for 15 min, and EU incorporation was detected using reagents from the Click-IT Plus EdU Flow Cytometry Assay following the EdU protocol.

### Nascent RNA sequencing

HT1080 cells were treated with 200 μg/ml bleomycin or 0.5 μM CBL0137 in triplicate. After 24 hr, cells were incubated with 1 mM EU for 20 min. RNA was isolated using the Monarch Total RNA Miniprep Kit (New England BioLabs, cat #T2010) following the manufacturer’s protocol. Nascent RNA was labeled with biotinylated azide and captured on magnetic beads using the Click-iT Nascent RNA Capture Kit (Molecular Probes, cat # C10365). cDNA was synthesized from RNA bound to the beads using the SuperScript VILO cDNA Synthesis Kit (ThermoFisher Scientific, cat #11754-050). cDNA was sequenced with the Illumina NextSeq 500/550 at the Roswell Park Comprehensive Cancer Center Genomics Shared Resource using Illumina NextSeq 500/550.

### Cell treatment for single-cell RNA sequencing

Populations of NDFs or HT1080 cells expanded after electroporation with approximately 50% of 53-positive and p53-negative cells were treated with 1 μM CBL0137 or 500 μg/ml bleomycin for 24 hr and then labeled with hashtag antibody oligonucleotide conjugate (HOT) from Biolegend (San Diego, CA) as follows: untreated cells, TotalSeq™-B0258 anti-human Hashtag 8 antibody; CBL0137-treated cells, TotalSeq™-B0259 anti-human Hashtag 9 antibody; bleomycin-treated cells, TotalSeq™-B0260 anti-human Hashtag 10 antibody. The different treatment groups for a given cell type were processed and sequenced together as one sample. Antibody binding and stability at the cell surface over several hours were tested by flow cytometry prior to the experiment.

### 10× Genomics

Single-cell libraries were generated using the 10× Genomics platform. Cell suspensions were assessed by trypan blue using the Countess FL automated cell counter (Thermo Fisher Scientific) to determine the concentration, viability, and absence of clumps and debris that could interfere with single-cell capture. Cells were loaded into the Chromium Controller (10× Genomics), where they were partitioned into nanoliter-scale gel beads-in-emulsion with a single barcode per cell. Reverse transcription was performed, and the resulting cDNA was amplified. The amplified full-length cDNA was used to generate gene expression libraries by enzymatic fragmentation, end-repair, a-tailing, adapter ligation, and PCR to add Illumina-compatible sequencing adapters. The resulting libraries were evaluated with D1000 ScreenTape using a TapeStation 4200 (Agilent Technologies) and quantitated using the Kapa Biosystems qPCR quantitation kit for Illumina. The libraries were pooled, denatured, and diluted to 300 pM with 1% PhiX control library added. The resulting pool was loaded into the appropriate NovaSeq Reagent cartridge and sequenced on a NovaSeq6000 following the manufacturer’s recommended protocol (Illumina Inc.).

### MNAse-sequencing

Micrococcal nuclease (MNase) was purchased from Worthington Biochemical Corp (Lakewood, NJ, Cat# LS004797). MNase digestion was done as described (28) followed by the analysis of fragment length by BioAnalyzer (Agilent Technologies, Inc., Santa Clara, CA). Digestion conditions resulting in predominantly monomucleosome fragments of 150-200 bp were used for DNA isolation. The sequencing libraries were prepared with the HyperPrep Kit (KAPA BIOSYSTEMS), from 1ug DNA. Following manufacturer’s instructions, the first step repairs the ends of the DNA fragments and a single ‘A’ nucleotide is then added to the 3’ ends of the blunt fragments. Indexing adapters, containing a single ‘T’ nucleotide on the 3’ end of the adapter, are ligated to the ends of the dsDNA, preparing them for hybridization onto a flow cell. Adapter ligated libraries are amplified by 3 cycles of PCR, purified using AMPureXP Beads (Beckman Coulter), and validated for appropriate size on a 4200 TapeStation D1000 Screentape (Agilent Technologies, Inc.). The DNA libraries are quantitated using KAPA Biosystems qPCR kit, and are pooled together in an equimolar fashion, following experimental design criteria. Each pool is denatured and diluted to 300pM with 1% PhiX control library added. The resulting pool is then loaded into the appropriate NovaSeq Reagent cartridge for 50 cycle paired-end reads and sequenced on a NovaSeq6000 following the manufacturer’s recommended protocol (Illumina Inc.).

### Bioinformatics analyses

Nascent RNA-seq paired-end raw sequencing reads passed quality filter from Illumina RTA were first pre-processed by using FastQC (v0.11.5) {Leggett, 2013 #1832} for sequencing base quality control. Reads were mapped to the Human Reference Genome (NCBI Build 37, GRCh37 (hg19)) using BWA (v0.7.15) {Li, 2010 #1833}. After marking PCR duplicates using Picard tool (v2.20.5, Broad Institute, Cambridge, MA) a second pass QC was done using alignment output with RSeQC (v2.6.3) {Wang, 2012 #1834} to examine abundances of genomic features and gene-body coverage. Gene expression is quantified according to hg19 RefSeq annotation using featureCounts from Subread aligner (v1.6.0) {Liao, 2013 #1835} with --fracOverlap 1 option. Differential expression analyses were performed using DESeq2 (v1.18.1) {Love, 2014 #1836}, a variance-analysis package developed to infer the statically significant difference in RNA-seq data. Downstream and visualization plots were done using regularized-log2 transformation implemented by DESeq2. For the scRNA-seq Chromium 10× Genomics libraries, raw sequencing data were processed using Cellranger software (http://software.10xgenomics.com/single-cell/overview/welcome) with GRCh37 (hg19) reference genome and GENCODE annotation. cloupe files generated by Cellranger was used with desktop Loupe Browser (10× Genomics) for cell filtering, analyses and visualization. First, data were filtered using the following thresholds: Unique Molecular Identifier (UMI) per barcode (cell) 30,000 – 130,000, features (genes) per barcode (cell) – 6,000 – 8,000, maximal mitochondrial counts – 7.5%. Then, cells were demultiplexed with hashtag oligos (HTOs) and assigned to the corresponding sample using threshold of Log2 >8 for of each antibody barcode count. Then, cells were assigned as p53 positive if *TP53* counts > 0 (linear) and *MDM2* counts log2 >1 and as p53 negative if *TP53* counts = 0 (linear) and *MDM2* counts log2 ≤1. Gene Set Enrichment Analysis was done using MSigDB online resource (Broad Institute, Cambridge, MA).

Raw GRO-seq reads from GSM1055806 were mapped to the Human Reference Genome (NCBI Build 38 (hg38)) using STAR (29). The resulting sequence alignment map (SAM) files were filtered using mapping quality (MAPQ > 10), sorted, and converted to the BAM format using samtools 1.10 (30). Nascent transcription profiles and metagene plots were generated using deepTools 2.0 (31). The pausing index was calculated as the ratio of the density of the reads near the TSS (+ 50 bp downstream) to the density of the reads in the whole gene.

Raw MNase-seq reads were mapped to the human genome hg38 using Bowtie2 (32) with parameter – very-sensitive. PCR duplicates in the alignments were filtered using the rmdup command from samtools 1.10. Sam files were sorted and converted to bam files using samtools 1.10. Bigwig files and metagene plots were generated using deepTools 2.0.

ChIP-seq and DNAse I sensitivity datasets for NDFs (ENCFF302JEV, ENCFF241ZSS, ENCFF087PPB, ENCFF715BVU, ENCFF969ORH, ENCFF471BTF, ENCFF429ZOO, ENCFF751GHM, ENCFF684QGK, and ENCFF087BQO) were downloaded from the ENCODE Project.

**Statistical analyses** were performed using GraphPad Prism software. The methods used for the analyses if individual experiments are indicated in the figure legends.

## Results

### Contrary to DNA damage, chromatin damage does not induce senescence

As expected, bleomycin, but not CBL0137, within 24 h of exposure caused DNA damage (DD), detected in the comet assay (Supplementary Figure S1A) and by staining for phosphorylated histone H2AX (γH2AX) (Supplementary Figure S1B), both markers of DNA breaks. In contrast, CBL0137, but not bleomycin, caused chromatin damage (CD) manifested as chromatin trapping (c-trapping) of histone chaperone FACT as the redistribution of FACT subunits, SSRP1 and SPT16, from the nucleoplasm to the chromatin (Supplementary Figure S1C) as a result of FACT binding to nucleosomes with unwrapped DNA (6).

We next compared the effects of DD and CD on the viability of two human mesenchymal cell types normal dermal fibroblasts (NDF) and human fibrosarcoma cell line HT1080. These cells have diploid genome and wild type p53. Both CD and DD decreased the viability of tumor cells more than normal cells (Figure 1A, B), although this relationship was reversed for bleomycin at lower concentrations (Figure 1B). Interestingly, the transition from no toxicity to full toxicity was much steeper for CBL0137 (occurring between 0.3 and 1 μM) than for bleomycin (toxicity gradually increased through three orders of magnitude from 1 to 1000 μg/ml) (Figure 1A, B and Supplementary Figure 2A, B). DD and CD suppressed DNA replication to a similar extent (Figure 1C).

**Figure 1.**
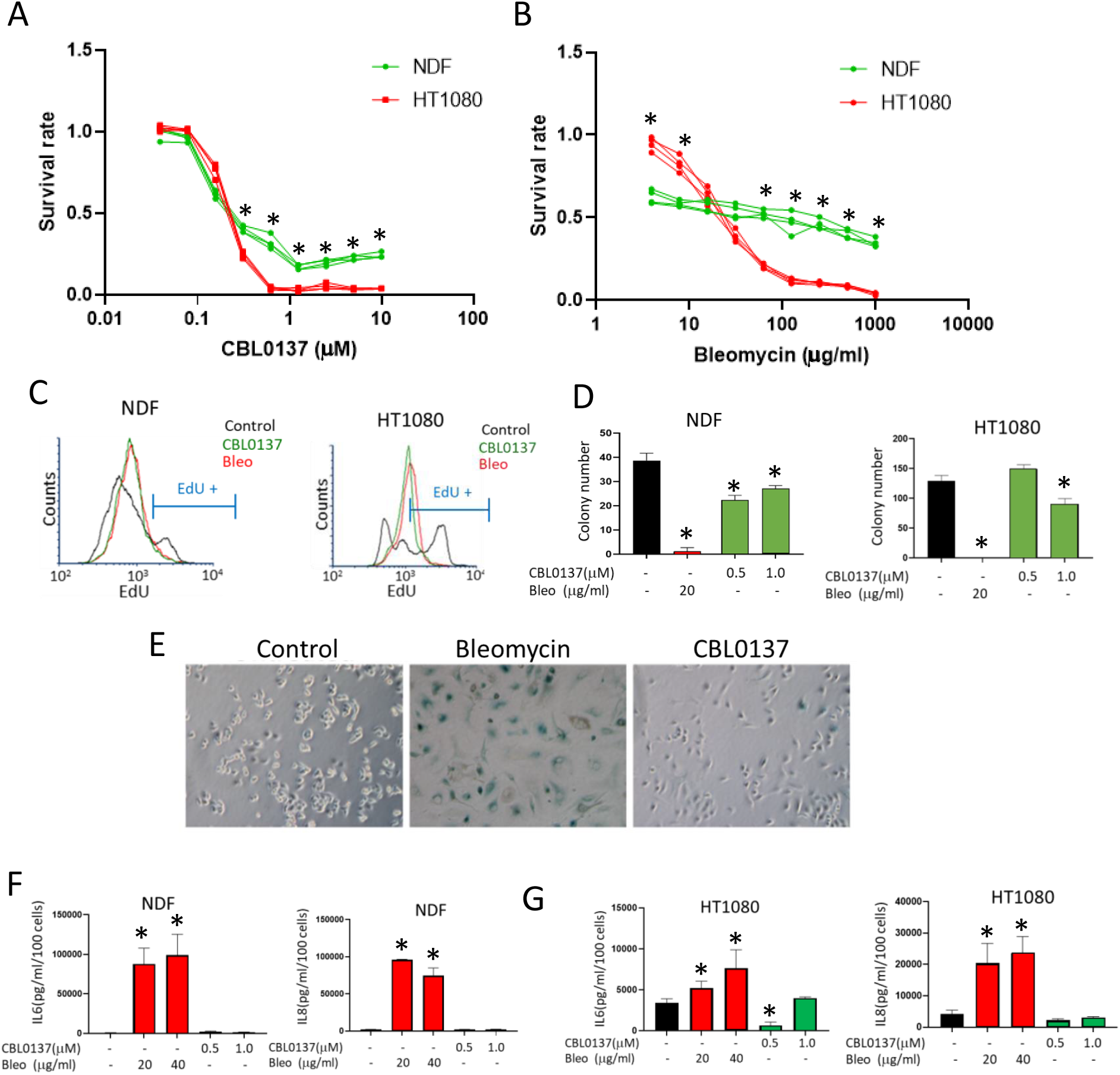
Comparison of the cellular effects of DD and CD. A, B. Cytotoxicity of CBL0137 and bleomycin in NDFs (A) and HT1080 cells (B) after 72 hours assessed by resazurin staining. The lines represent four replicates of one representative experiment normalized to untreated cells. *p < 0.05 by the Student’s t-test, NDFs vs. HT1080 cells. C. EdU incorporation assay comparing replication in NDFs and HT1080 cells treated with 500 μg/mL bleomycin (Bleo) or 1 μM CBL0137 for 24 hours. EdU was added 2 hours before the cells were fixed and stained. Histograms show the distribution of EdU-positive cells in the different samples. The marker shows the area of untreated EdU-positive cells. D-G. Assessment of the senescence phenotype in cells treated with bleomycin or CBL0137 for 24 hours followed by 10 days in drug-free medium. After 10 days, the cells were split and assessed for ability to proliferate (colony formation - D), acidic beta-galactosidase staining (E), and secretion of SASP factors, IL6 (F) or IL8 (G). D, F, and G. Data are presented as the mean ± SD (n = 3). *p < 0.05 by the Student’s t-test vs. untreated cells.

DD and CD differed in their ability to cause senescence. Bleomycin is the most frequently used chemical to induce senescence *in vitro* and *in vivo* (33). Consistent with these findings, we observed the emergence of senescent markers in cells treated with bleomycin dose as low as 20 μg/ml. In contrast, low CBL0137 doses did not affect cell growth, whereas higher doses caused cell death within 48 hr. At a narrow range of concentrations (0.5–1 μM) and limited time of incubation (up to 24 hours), CBL0137 caused growth arrest; however, these cells were able to proliferate and form colonies from single cells after the compound was washed out (Figure 1D). Cells arrested by CBL0137 did not develop markers typical for senescent cells, such as positive staining for acidic beta-galactosidase (Figure 1E) or senescence-associated secretory phenotype (Figure 1F, G). Thus, major difference between DD and CD in their cytotoxic effects was inability of CD to induce senescence.

### Different dynamics of p53 activation by DD and CD

An important common feature of DD and CD is the activation of the p53 pathway (5,34). However, we found that the dynamics of p53 protein accumulation differed between DD and CD. For example, a wide range of bleomycin concentrations used caused similar elevations in p53 protein levels, whereas CBL0137 induced a sharp increase in p53 protein with a peak between 0.6 μM and 1.2 μM followed by a decrease (Figure 2A, B). The kinetics of the p53 protein increase was similar between the two compounds and cell types (Figure 2C). Interestingly, higher p53 levels accumulated in response to CD than to DD in both cell types. However, the p53 accumulated in response to CD lacked posttranslational modifications typical for DD, such as N-terminal phosphorylation and Lys 382 acetylation (5).

**Figure 2.**
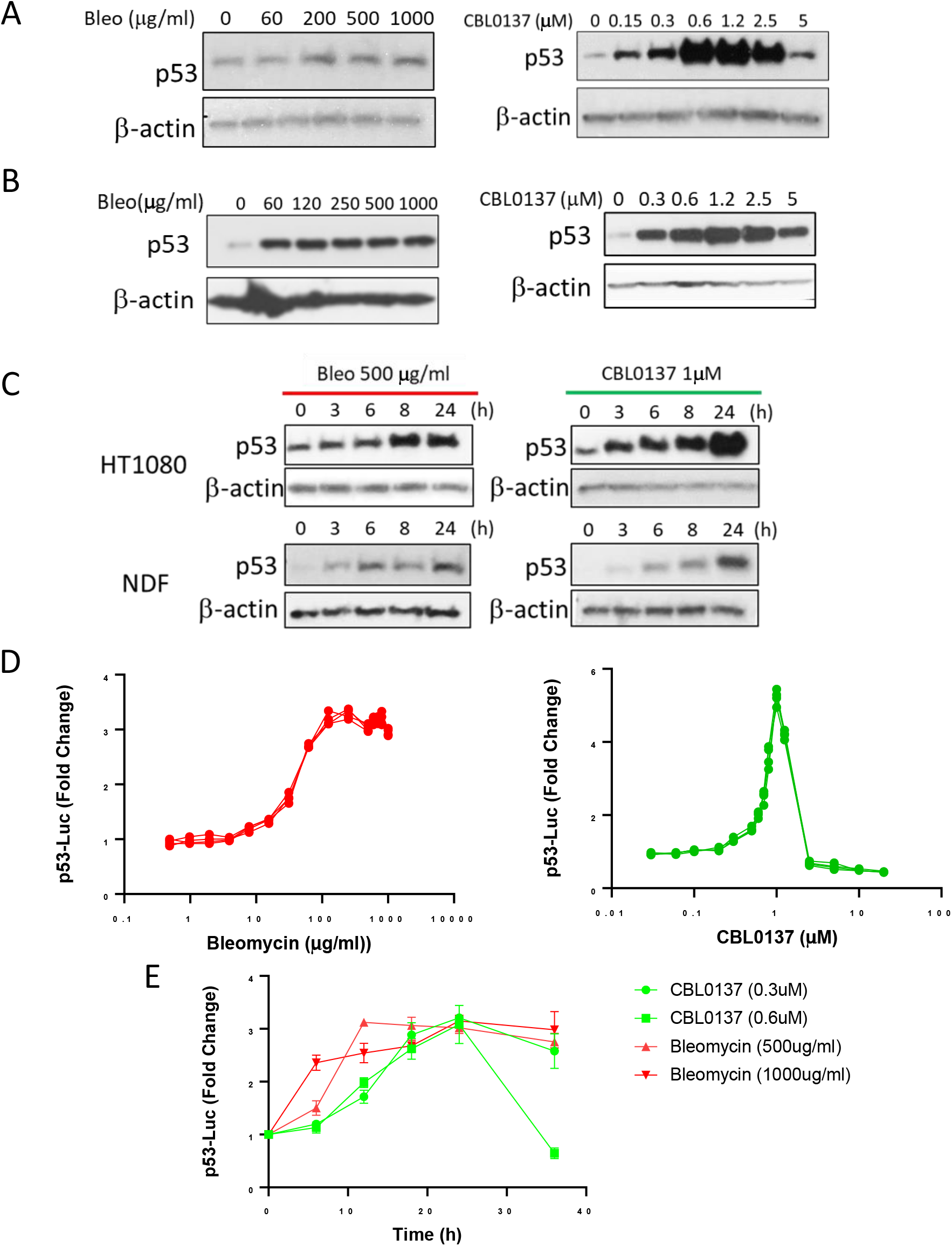
p53 activation by DD and CD. A, B. Western blotting of lysates from NDFs (A) and HT1080 cells (B) treated with different concentrations of bleomycin (bleo) or CBL0137 for 24 hours. C. Western blotting of lysates from NDFs and HT1080 cells treated with 500 μg/ml bleo or 1 μM CBL0137 for the indicated times. D, E. Induction of p53-responsive luciferase reporter in HT1080 cells treated with different concentrations of the drugs for 24 hours (D) or the indicated concentrations for different periods of time (E). In (D), three replicates for each drug are shown. In (E), data are presented as the mean ± SD (n = 3 replicates).

To confirm that unmodified p53 accumulated upon CD is transcriptionally active, we used HT1080 cells with an integrated luciferase reporter controlled by a p53 consensus element and p53 binding site from the *CDKN1A* promoter (22,35). In line with p53 protein accumulation, p53 transcriptional activity had different dynamics in the presence of DD and CD agents. The p53-responsive transcription quickly reached a plateau and remained at the same level over a wide range of bleomycin concentrations. The CBL0137 dose-response curve for p53 transcriptional activity was bell-shaped with a narrow peak at approximately 0.5–1.5 μM (Figure 2D). The kinetics for p53 transactivation paralleled that of p53 accumulation for both agents (Figure 2E). Thus, DD caused less intensive but steady state p53 activation at a wide concentration range, whereas p53 accumulation in response to CD was higher but limited to a narrow concentration range.

### CD induces broader changes in gene expression than DD

To compare the transcriptional response of cells to DD and CD, we selected concentrations of both agents that caused maximal p53 accumulation and reporter activation 24 hours after the start of treatment (500 μg/ml bleomycin and 1 μM CBL0137). To separate p53-dependent and -independent changes in gene expression, we deleted the p53 gene from HT1080 cells and NDFs using the CRISPR/Cas9 approach. To avoid potential DD from constitutive Cas9 expression in cells, we electroporated cells with recombinant Cas9 protein and reconstituted the RNP complex, consisting of four synthetic gRNAs targeting the first and the last exons of p53. Normally, this approach requires cell cloning and sequencing of target regions to identify the biallelic deletion. However, we used single-cell RNA sequencing (scRNA-seq) of the whole cell population after electroporation to avoid potential artifacts associated with clonal variability. This method allowed us to distinguish cells with and without functional p53 by their transcriptome, and we determined the proportion of cells that lost p53 after electroporation by p53 staining (Supplementary Figure S3). Subsequent experiments were performed with cell populations consisting of an approximately 1:1 ratio of p53 negative and p53 positive cells. Labeling cells treated under different conditions (control, CBL0137, and bleomycin) with different oligonucleotide-conjugated antibodies allowed library preparation and sequencing of all samples simultaneously as one pool for each cell type.

Unbiased clustering and UMAP plots demonstrated that both normal and tumor cells formed a diagonal axis along UMAP1 and UMAP2 with groups of control and CBL0137-treated cells located most distantly from each other and a group of bleomycin-treated cells located between those two groups and closer to the control (Figures 3A and 4A), suggesting that CD has stronger effects on the cell transcriptome than DD.

**Figure 3.**
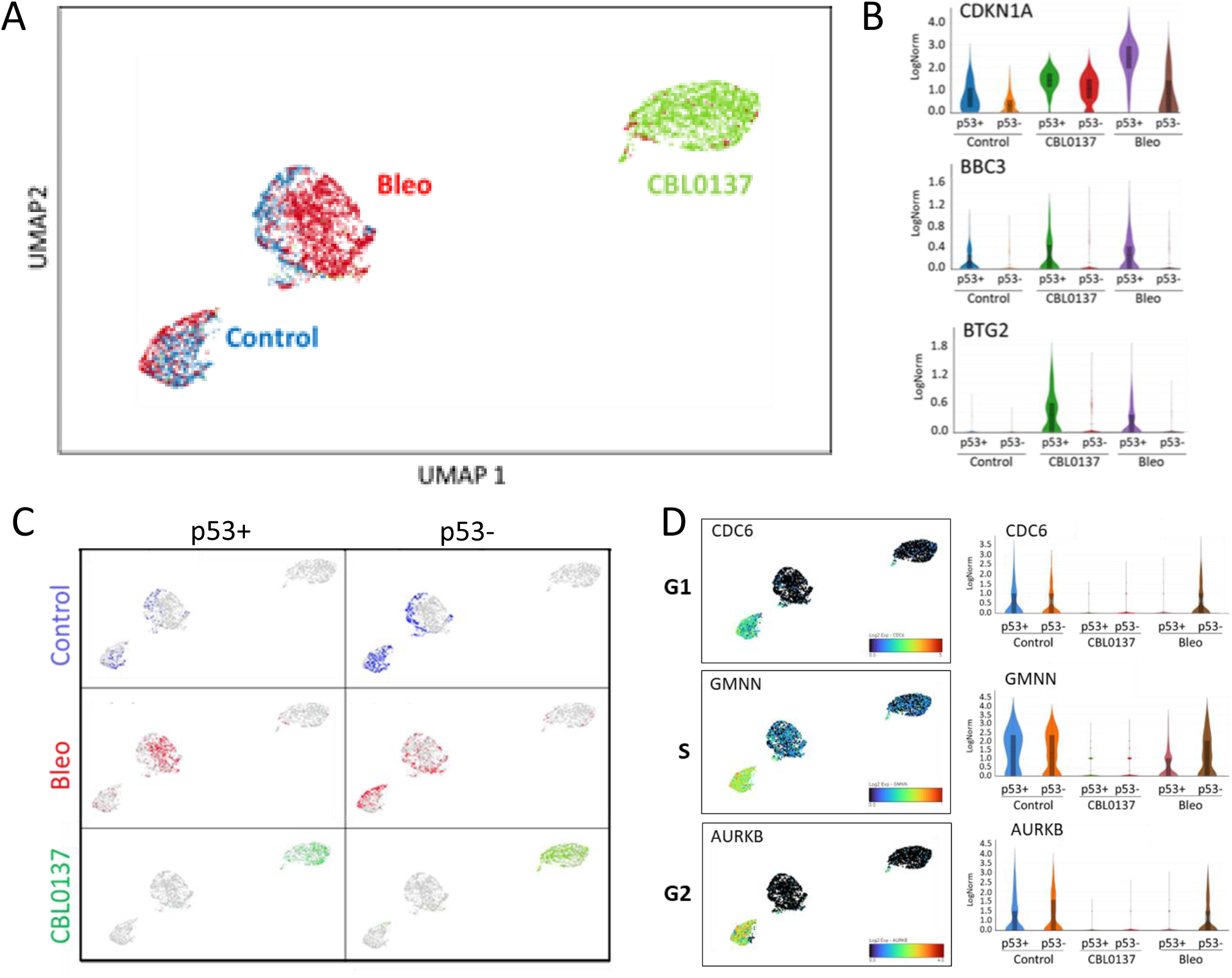
Effects of CD and DD on cell transcriptome of NDF cells detected by scRNA-seq. A. UMAP plot shows the positions of– control (blue), bleomycin (bleo)-treated (red), or CBL0137-treated (green) cells. B. Violin plots showing the distribution of expression of p53 regulated genes under different conditions and between cells classified as p53-positive or p53-negative. C. UMAP plots of NDF cells color-coded based on the treatment condition (as in A) and p53 status. D. UMAP and violin plots showing the expression of markers for the G1, S, and G2/M phases of the cell cycle in NDFs cells under different conditions.

**Figure 4.**
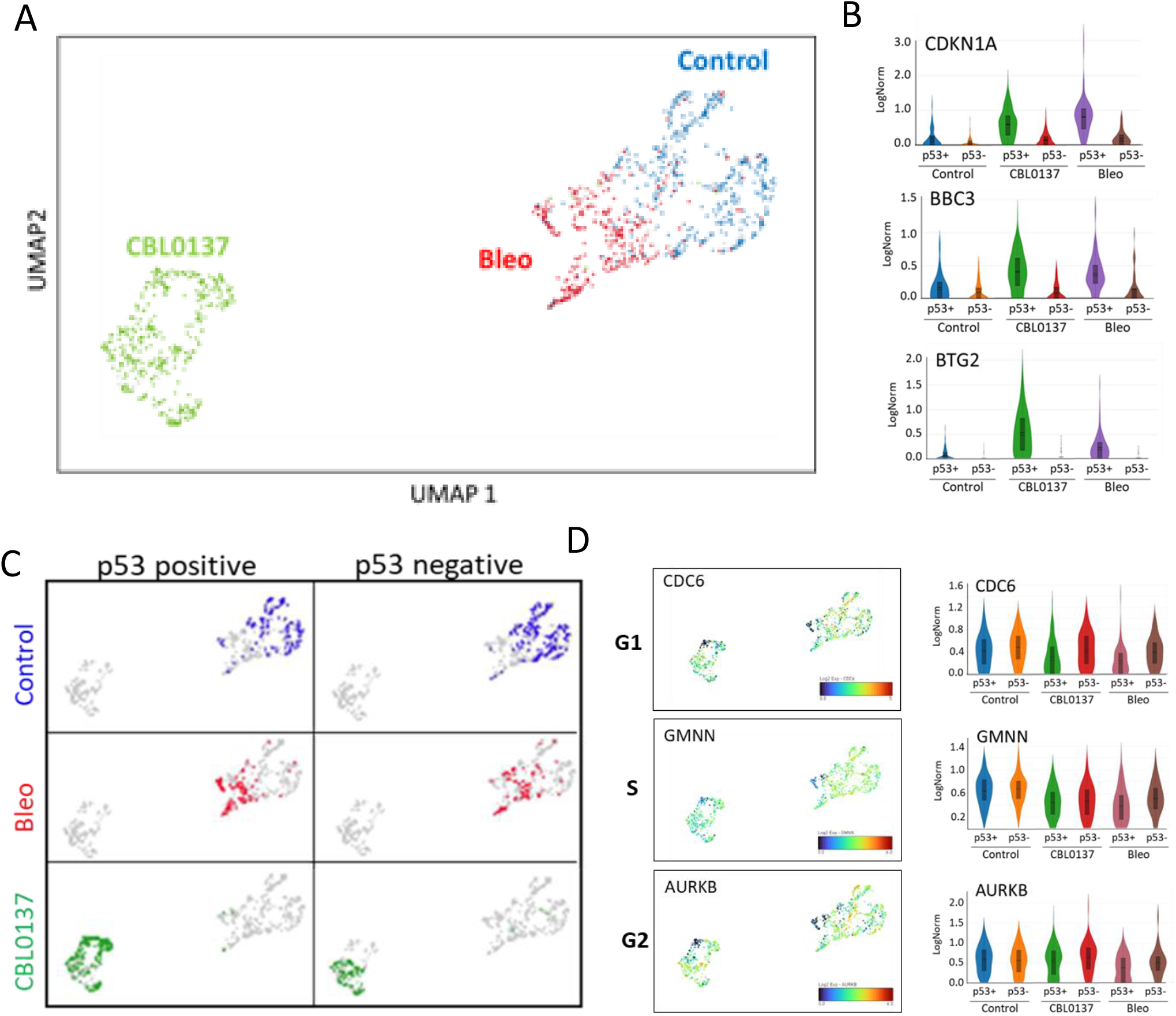
Effects of CD and DD on cell transcriptome of HT1080 cells detected by scRNA-seq. A. UMAP plot shows the positions of– control (blue), bleomycin (bleo)-treated (red), or CBL0137-treated (green) cells. B. Violin plots showing the distribution of expression of p53 regulated genes under different conditions and between cells classified as p53-positive or p53-negative. C. UMAP plots of HT1080 cells color-coded based on the treatment condition (as in A) and p53 status. D. UMAP and violin plots showing the expression of markers for the G1, S, and G2/M phases ofthe cell cycle in HT1080 cells underdifferent conditions.

To identify p53-positive and p53-negative cells, we first looked for the presence of reads corresponding to *TP53;* however, most cells had very low numbers of TP53 reads (0-3) independent of cell type or treatment. Therefore, we also used reads corresponding to the most specific p53 target, *MDM2* (Supplementary Figure S4A-H, details are in Materials and Methods) to distinguish p53 positive and negative cells. Other p53 targets showed expression patterns similar to MDM2 (Figures 3B and 4B). Surprisingly, this analysis showed that p53-positive and p53-negative cells were located in the same overlapping regions of the UMAP plots for the control and CBL0137-treated cells or very close in the case of bleomycin-treated cells (Figures 3C and 4C). Thus, although p53-positive and p53-negative cells could be distinguished by the expression of several p53 target genes, the p53 status had a weak impact on the cell transcriptome in response to DD and almost no impact in response to CD or under unstressed conditions.

For the NDFs, untreated and bleomycin-treated cells were located very close to each other on the UMAP plots, although they did not overlap (Figure 3A). These cells were found within two clusters: one mostly consisting of untreated cells (Figure 3A, left) with a small admixture of bleomycin-treated cells, and the second cluster was comprised predominantly of bleomycin-treated cells with an “edge” of untreated cells (Figure 3A, center). A major difference between these two clusters was in the expression of cell cycle markers, including genes predominantly transcribed during G1 (*CDC6*, *CDC25B*), S (*CDC45, MCM4, EXO1, GMMN1*), and G2/M (*AURKB, MKI67*) phases of cell cycle (Figure 3D, Supplementary Figure S4I). The left cluster was positive for these markers and consisted of control cells and p53-negative bleomycin-treated cells. The central cluster negative for the cell cycle markers contained p53-positive bleomycin-treated cells and non-cycling control cells (Figures 3A, C, Supplementary Figure S4I). These results suggest that the major response of normal cells to DD is p53-dependent growth arrest. Moreover, p53-positive bleomycin-treated cells were not different from non-cycling control cells, reinforcing the concept that normal cells respond to DD with growth arrest.

Control and bleomycin treated HT1080 cells occupied two adjacent regions within one cluster (Figure 4A). The cells within this cluster formed a gradient transition from high to low expression of cell cycle markers and from low to high p53 positivity without clear boundaries between samples (Figure 4A, 4D and Supplementary Figure S4F, G, H, J). This gradient started from the untreated cells (the highest cell cycle marker levels) followed by p53-negative bleomycin-treated cells (high cell cycle markers) and then p53-positive bleomycin-treated cells (lower cell cycle marker levels). The difference between the expression levels of the cell cycle markers was less pronounced in tumor than normal cells (Compare figure 3D vs.4D and Supplementary Figure S4I vs.S4J), suggesting that in contrast to normal cells, there is more variability in the tumor cell response to DD with a less pronounced cell cycle arrest and less dependence on p53.

CBL0137-treated cells were clearly separated from the other cell groups (Figures 3A and 4A), consistent with many more genes which expression was changed in response to CD than DD: CD altered the expression of 1375 genes in p53-positive NDFs, 1304 genes in p53-negative NDFs, 543 genes in p53-positive HT1080 cells, and 309 genes in p53-negative HT1080 cells (p_adj_ < 0.05) (Figure 5A). In contrast, DD changed the expression of only 107 genes in p53-positive NDFs, 4 genes in p53-negative NDFs, 117 genes in p53-positive HT1080 cells, and 2 genes in p53-negative HT1080 cells (Figure 5A). Thus, CD caused much more dramatic changes in the cell transcriptome than DD. Moreover, there was a striking p53 dependence in response to DD, whereas CD caused a transcriptional response with a magnitude indicative of a p53-independent response (Figure 5B, C). Furthermore, most genes whose expression changed in response to CD were the same in the p53-positive and p53-negative cells (Figure 5D–G), further supporting the hypothesis that the transcriptional response to CD is largely p53-independent.

**Figure 5.**
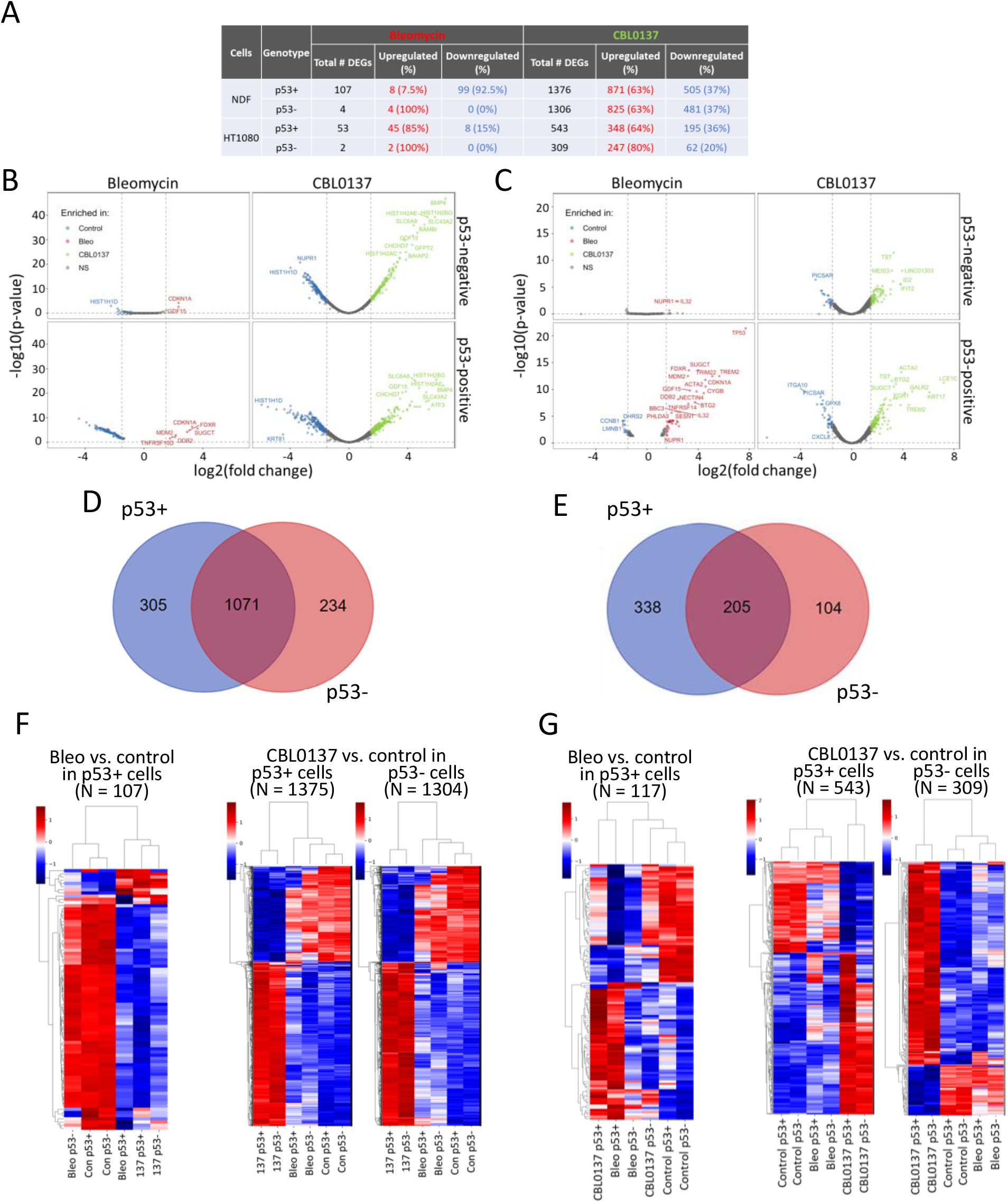
Transcriptional response to CD is p53-independent. A. Number of differentially expressed genes (DEGs, FC > 1.5, adjusted p-value < 0.05) between different treatment conditions in p53-positive and p53-negative cells. B, C. Volcano plots showing the DEGs in p53-positive and p53-negative NDFs (B) and HT1080 cells (C) under different treatment conditions (control [Con], blue; bleomycin [Bleo], red; CBL0137, green). D, E. Venn diagram showing the overlap of DEGs in p53-positive and p53-negative NDFs (D) and HT1080 cells (E). F, G. Dendrograms and heatmaps showing unbiased hieratical clustering of samples and genes in NDFs (F) and HT1080 cells (G). The number of DEGs (N) with padj < 0.05 is indicated at the top of each plot. No plots were built from the DEGs of p53-negative bleomycin-treated cells due to the low number of DEGs (see panel A).

### Gene expression changes in response to DD reflect p53-dependent growth arrest

The transcriptional response to DD was strikingly different between the two p53 genotypes. Although the expression levels of a substantial number of genes were changed in the p53-positive NDFs (107 genes) and HT1080 cells (53 genes) following treatment with bleomycin, the expression of only 4 genes in the NDFs and 2 genes in the HT1080 cells were changed in response to DD in p53-negative cells, demonstrating the major role of p53 in the DD response (Figure 5A–C). Only 8 out of 107 changed genes were upregulated in the NDFs, among which four were p53 targets (*CDKN1A, MDM2, DDB2*, and *FDXR*). The 99 suppressed genes were highly enriched for ‘E2F targets’ and ‘G2M checkpoint’ (Figure 6A) and represented almost exclusively genes expressed during the cell cycle, indicating these cells were growth arrested. The ratio between the upregulated and downregulated genes was reversed for the tumor cells: 45 genes were upregulated, and eight were downregulated. The upregulated genes were strongly enriched for ‘p53 targets’ (18 out of 45), whereas the downregulated genes were enriched for ‘E2F targets’ and ‘G2M checkpoint’ (6 of 8 combined) (Figure 6B). Thus, the response to DD in these cells was almost exclusively through p53 and included p53-dependent growth arrest, which was stronger in normal cells than in tumor cells.

**Figure 6.**
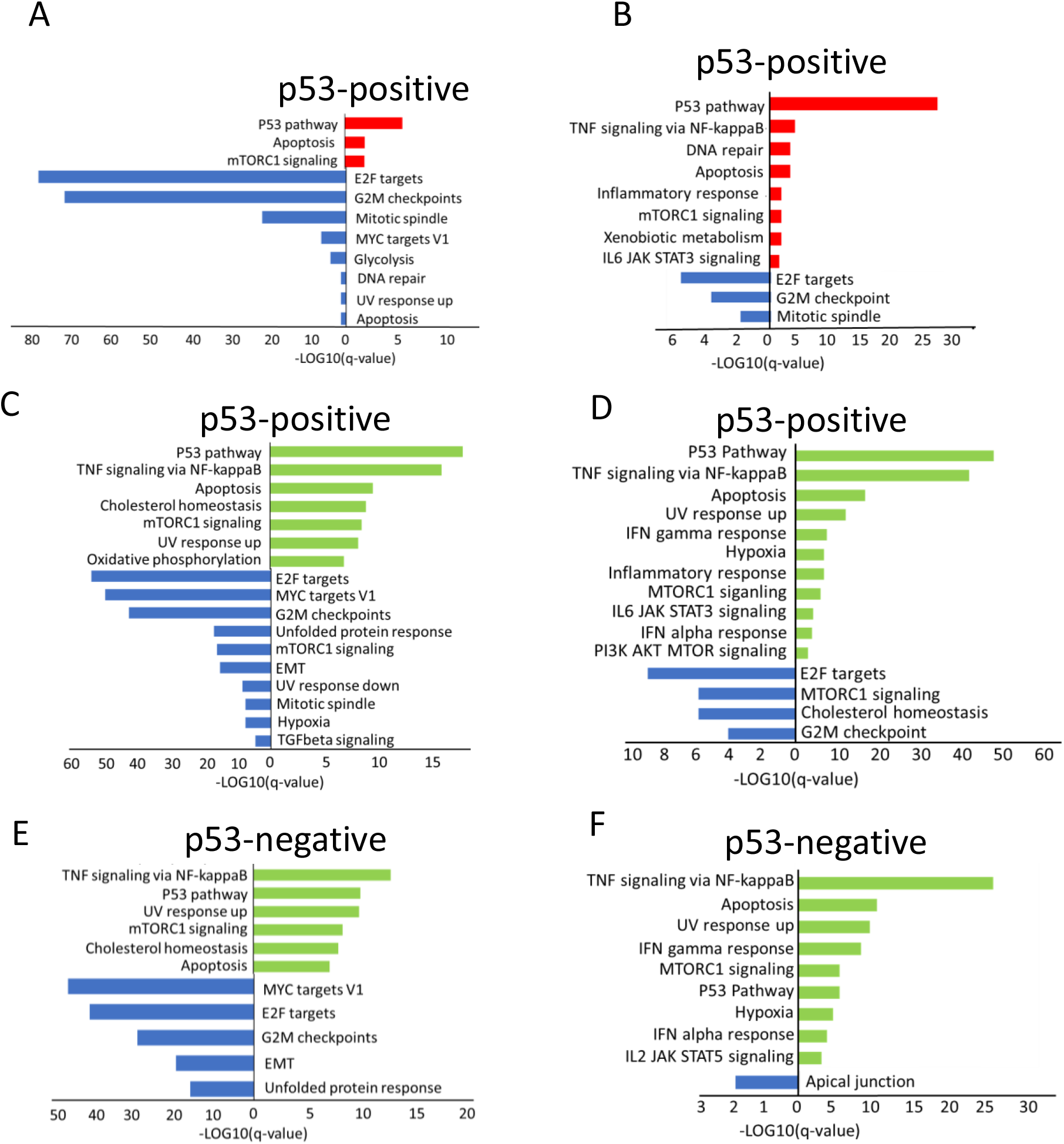
Gene set enrichment analysis of differentially expressed genes between different conditions identified by scRNA-seq in NDFs (A, C, E) and HT1080 cells (B, D, F). A, B. Hallmark gene sets enriched in control (blue) or bleomycin-treated (red) p53-positive cells. C, D. Hallmark gene sets enriched in control (blue) or CBL0137-treated (green) p53-positive cells. E, F. Hallmark gene sets enriched in control (blue) or CBL0137-treated (green) p53-negative cells.

### Minimal effect of p53 on the transcriptome of control and CBL0137-treated cells

To look more carefully at the influence of p53 on transcription, we compared gene expression in p53-positive and p53-negative cells under different conditions. Only four genes in the NDFs (*TP53, MDM2, CDKN1A*, and *GDF15*) and seven in the HT1080 cells (*TP53, MDM2, CDKN1A, GDF15, FN1, ZMAT3*, and *FDXR*) two of which were used for cell classification, varied between untreated p53-positive and p53-negative cells. The expression levels of these genes were all higher in p53-positive cells. This minimal changes suggested that p53 had a minimal effect on gene expression in unstressed cells in line with very low mRNA and protein levels. Surprisingly, a similar situation was observed in CBL0137-treated cells: 12 genes in the NDFs and 43 genes in the HT1080 cells were significantly higher in p53-positive cells versus negative cells, and no genes were significantly lower. Although there were p53 targets among the genes expressed higher in CBL0137-treated p53-positive cells than in CBL0137-treated p53-negative cells (3 out of 12 genes in NDF and 20 out of 43 in HT1080), this low number of genes differing between genotypes relative to the total number of differentially expressed genes resulting from CD (Figure 5A), confirms that the CD response is primarily p53-independent.

There were consistently more genes upregulated than downregulated in response to CD in both genotypes and cell types (Figure 5). Upregulated genes mainly belonged to two lists, ‘p53 targets’ and ‘NF-kappaB activation via TNF’. These lists were enriched within both genotypes (Figure 6A-F). Genes downregulated upon CD, similarly to DD were enriched for ‘E2F targets’ and the ‘G2M checkpoint’ (Figure 6C–F).

To confirm that the CD response is largely p53-independent, we used additional methods of p53 inactivation and assessment of transcription. We inactivated p53 in HT1080 cells using the dominant-negative mutant GSE56 that prevents p53 oligomerization (23) and then sequenced newly synthesized transcripts labeled with EU for 20 min at the end of the 24-hr treatment. As expected, bleomycin did not induce p53 targets in the cells with inactive p53; however, there was almost no difference in the responses of p53 target genes to CBL0137 between cells with active or inactive p53 (Supplementary Figure S5).

Our experiments clearly demonstrated different changes in the cell transcriptome in response to DD and CD. There were many more genes which expression was changed in response to CD than DD. While response to DD was highly p53 dependent, response to CD was largely p53 independent. At the same time, CD induced the expression of p53-dependent genes. The biggest puzzle is that these “p53-dependent” genes were also induced in p53-negative cells.

### Genes activated by CD have a unique chromatin state at their promoters

P53 targets were the most enriched category among genes activated by CD in all cells (i.e., normal and tumor, p53-positive and p53-negative). Besides being enriched within the category of ‘p53 target genes’, these genes were also enriched within the ‘NF-kappaB targes’ and several other categories that collectively could be characterized as ‘stress-response genes’ (Figure 6 C–F). Several reports have shown that many stress-response genes are regulated at the level of RNA polymerase II pausing (36,37). Under basal conditions, these genes have preassembled RNA polymerase II on their promoters engaged in abortive elongation until a specific transcription factor binds and releases RNA polymerase II into productive elongation through a currently undefined mechanism (38). Thus, we hypothesized that stress-response genes induced by CBL0137 might have a specific chromatin state around their TSS, making them sensitive to CD. Thus, we examined the state of chromatin around the TSS of genes activated by CD in both p53-positive and p53-negative cells under basal conditions (i.e., untreated) using our data and publicly available datasets. More data was available for NDFs, although the limited data for HT1080 cells confirmed our finding in the NDFs (Supplementary Figure S7A, C). We compared genes induced by CD with all other genes classified by their expression levels under basal conditions based on bulk (NDFs) or nascent (HT1080) RNA sequencing. Genes were divided into four quartiles: (i) no expression, (ii) low expression, (iii) moderate expression, and (iv) high expression levels. Based on the average expression of CD-induced genes under basal conditions, the CD-induced genes were close to the moderately expressed genes. Analysis of the publicly available GRO-seq data for IMR90 cells (human lung fibroblasts, (39)) confirmed this classification (Figure 7A, B).

**Figure 7.**
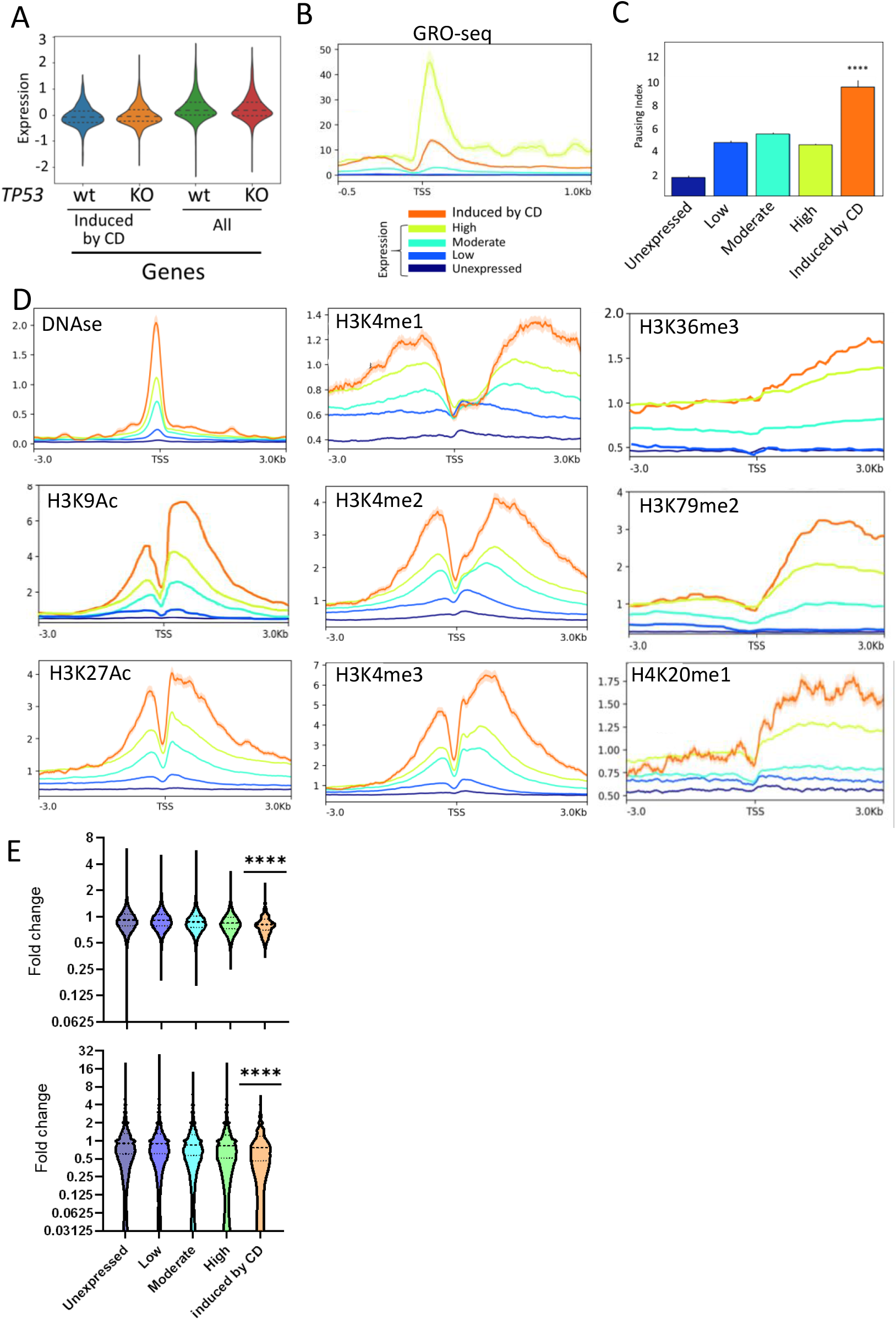
Genes activated by CD have a specific chromatin state around TSS. A. Comparison of expression levels of genes activated by CBL0137 in p53-positive and p53-negative NDFs with the expression levels of all other genes. Violin plots of normalized expression levels obtained by bulk RNA seq. B. Normalized profiles of GRO-seq for different categories of genes in NDFs. C. RNA polymerase II pausing index calculated using the GRO-seq data of IMR90 cells. ****p < 0.0001 by the Kruskal-Wallis test. D. Metagene profiles of DNAse hypersensitivity (DNAse) and the indicated histone post-translational modifications for different categories of genes. The same legend as in panel B. E. Change in MNase-seq read density at the indicated distances around the TSS of different genes in HT1080 cells treated with 1 μM CBL0137 for 1 hour versus the control. ****p < 0.0001 by two tailed t-test, CD-induced genes vs. other groups (comparison between other groups is not shown).

We looked at RNA polymerase II pausing in these quartiles using the IMR90 GRO-seq dataset. Consistent with our hypothesis, CD-induced genes had the highest pausing index in basal conditions (Figure 7C). Next, we evaluated DNAse hypersensitivity and histone marks associated with active and repressed states. DNA around the TSS of CD-induced genes was more sensitive to nuclease digestion than DNA of the most highly expressed genes (Figure 7D), suggesting the presence of nucleosome-free regions. Moreover, the nucleosomes present around the TSS of these genes had the highest levels of activating histone marks, including H3K27Ac, H3K9Ac, and mono-, di-, and trimethylated H3K4 (Figure 7D). Interestingly, these genes also had the highest levels of histone marks associated with active elongation downstream of their TSS, such as H3K36me3, H3K79me3, and H4K20me1 (Figure 7D). Repressive histone marks on CD-induced genes were as low as these marks at highly expressed genes (Supplementary Figure S6B).

These analyses showed that genes activated by CD have a specific chromatin state with a high level of paused RNA polymerase II. CD may activate transcription of these genes either because, in this chromatin state, nucleosomes are more sensitive to CD than anywhere else or because CBL0137 triggers other mechanisms that allow the release of paused RNA polymerase II. To distinguish between these hypotheses, we measured nucleosome loss from the same categories of genes upon treatment with CBL0137 using micronuclease (MNase) digestion followed by sequencing (MNase-seq) (28). Indeed, we found that on average CD induced gene lost more nucleosomes at and around TSS than genes from other categories (Figure 7E and Supplementary Figure S7C).

Although the genes activated by CD did not fall within the quartile of the most highly expressed genes, they had a chromatin composition at and around their TSS that was more open and more decorated with the histone marks for active transcription than the most highly expressed genes. Their gene bodies contained more histone marks of active elongation than the most actively transcribed genes and were enriched within different stress-response categories and had the highest RNA polymerase II pausing index. Nucleosomes at regions around TSS of these genes are also the most sensitive to nucleosome destabilizing effect of CD.

### Chromatin damage activates a p53-regulated reporter in the absence of p53

Higher nucleosome loss at promoters of CD induced genes suggests that CD may trigger transcription of these genes in the absence of specific transcriptional factor though direct effect on chromatin. This hypothesis is in line with similar degree of activation of p53 responsive genes in p53 positive and negative cells. Thus we decided to test whether CD can induce activity of very specific p53 reporter in the absence of p53. First, we excluded that CBL0137 activated general transcription using the EU incorporation assay (Figure 8A). Then we transduced a p53-responsive luciferase reporter construct into p53-positive and p53-negative HT1080 cells (Figure 8B). In untreated condition, reporter activity was > 500 times higher in the p53-wild type than in the knockout cells (Figure 8C). This was surprising, taking into account the very low number of p53 transcripts in cells and no major impact of p53 on the cell transcriptome under basal conditions. This suggests that just a few p53 molecules in some cells are enough to cause a transcriptional burst; however, this burst involves only a few genes and does not significantly affect cell transcriptional programs.

**Figure 8.**
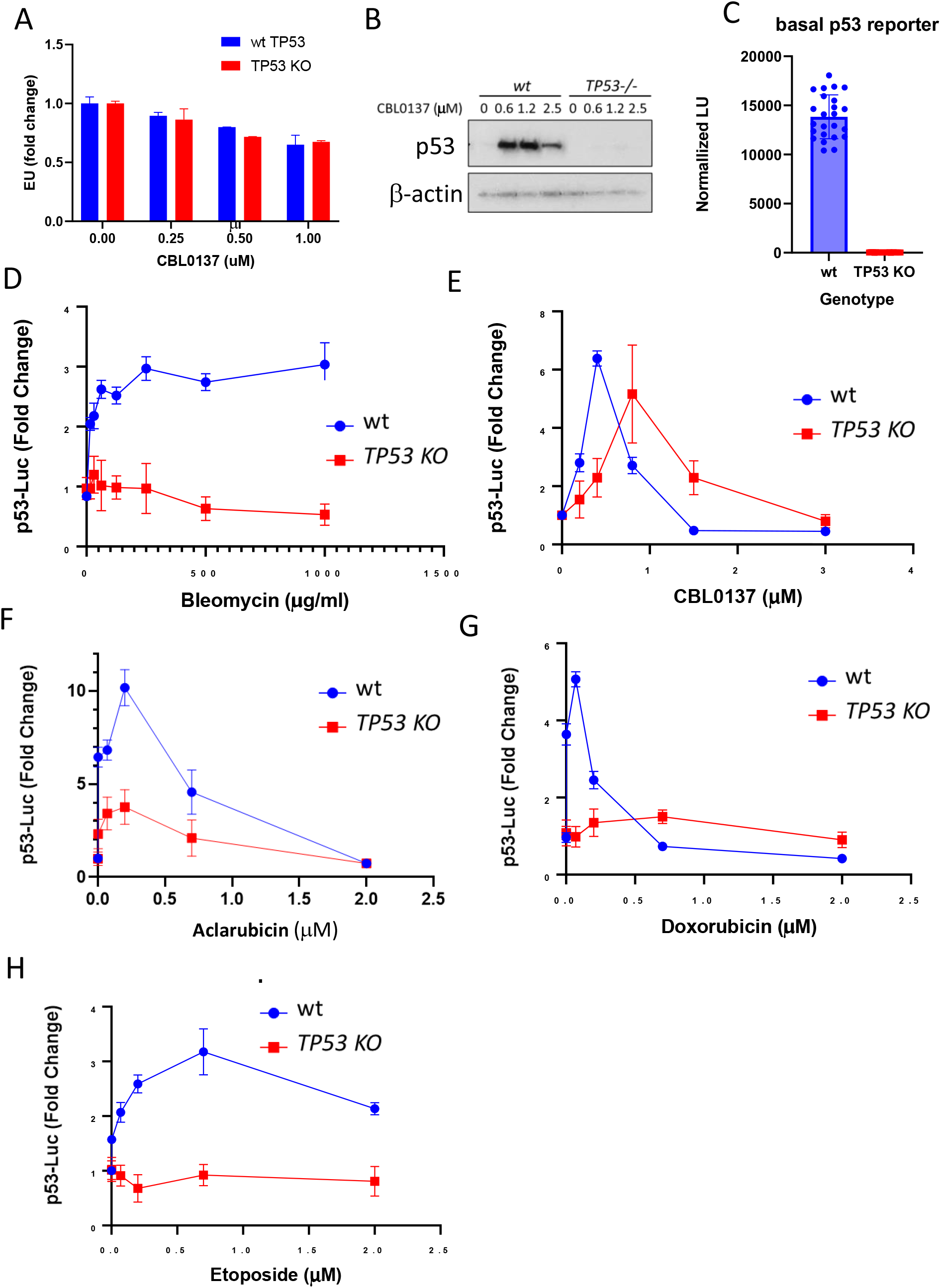
CD, but not DD, induces p53 transcriptional activity in the absence of p53. A. Effect of CBL0137 on general transcription. EU incorporation assay for p53-positive and p53-negative HT1080 cells treated with CBL0137 for 3 hours. Data are presented as the mean ± SD (n = 2 replicates). B. Western blotting of lysates from p53-positive and p53-negative HT1080 cells treated with CBL0137 for 24 hours. C. Basal luciferase activity controlled by a p53-responsive promoter in p53-positive and p53-negative HT1080 cells, normalized by cell number. Data are presented as the mean ± SD (n = 6 replicates in 3 experiments). D-H. Fold-change in p53-responsive luciferase activity in p53-positive and p53-negative HT1080 cells treated with bleomycin (D), CBL0137 (E), aclarubicin (F), doxorubicin (G), or etoposide (H) for 24 hours.

When we treated this pair of reporter cells with bleomycin or CBL0137 for 24 hours, we observed that DD activated the p53-responsive reporter in the p53 wild-type cells but had no effect on the reporter in p53-negative cells (Figure 8D). In contrast, CD activated the reporter regardless of p53 status, demonstrating that CD could activate transcription without a specific transcription factor (Figure 8E). Because this phenomenon was highly unusual, we tested other chemicals in this pair of reporter cell lines differing in their ability to cause CD or DD – two anthracyclines (doxorubicin and aclarubicin) and the podophyllotoxin etoposide. The two anthracyclines differ in their ability to cause DD or CD. Doxorubicin causes DD by inhibiting topoisomerase II and generating reactive oxygen species (40). It causes rather weak CD in cells (27). Aclarubicin causes less DD than doxorubicin but induces CD comparable to CBL0137 (27). Etoposide, a non-DNA-binding topoisomerase II inhibitor, causes DD but not CD (27). Both anthracyclines activated the p53 reporter in the absence of p53; the degree of activation corresponded their CD activity (Figure 8F, G). Like bleomycin, etoposide did not activate the p53-responsive reporter in the p53-negative cells (Figure 8H). Thus, our data suggest that although DD induces p53 targets by activating p53 protein, CD can activate these targets without functional p53 protein.

## Discussion

In this study, we identified the commonalities and differences in the response of normal and tumor cells to DD and CD. Most of our observations were performed shortly after the induction of DD by a drug mimicking nuclease activity or CD by a drug directly interfering with DNA-histone binding. We selected bleomycin because short-term treatment does not damage nucleosomal DNA, leaving nucleosomes intact (15,17,20,41), and CBL0137 because it does not cause detectable DNA damage ((5,11)). However, we cannot exclude CD as a later secondary effect of bleomycin because DNA breaks facilitate spontaneous DNA unwrapping from the histone core and chromatin disassembly during DNA repair. Furthermore, DD may be a secondary effect of CBL0137 because DNA unwrapped from the histone core is more exposed to water molecules, increasing the chances of encountering reactive oxygen species.

DD and CD are both cytotoxic; tumor cells are more sensitive to both types of damage than normal cells. Both types of damage also activate p53. However, both these consequences of DD and CD have features specific to the type of damage. The major difference in cytotoxicity induced by DD and CD is that DD causes cellular senescence, while CD does not. This difference likely explains the recently reported absence of cardiotoxicity from aclarubicin treatment, which causes predominantly CD (27). Doxorubicin is structurally similar to aclarubicin but primarily causes DD and has a strong cardiotoxic effect (18,19). Importantly, anti-cancer activity of both compounds were similar, suggesting that anti-cancer efficacy depends on CD, while the toxicity on DD.

Another interesting feature that distinguishes the toxic effects of DD and CD is that bleomycin toxicity is almost linearly proportional to the dose administered and possibly the number of DNA breaks, although we did not measure this. In contrast, CBL0137 shifts from non-toxic to fully toxic at a narrow concentration range. This phenomenon may be explained by one of two models: (i) chromatin tolerates the intercalation of small molecules into DNA without destabilizing nucleosomes up to a certain threshold, e.g., if the intercalator first binds to linker DNA; (ii) chromatin is gradually decondensed but cell viability is compromised only after a certain level of decondensation is reached. Mechanisms of toxicity attributed to chromatin decondensation are still obscure; one possible explanation is that free histones evicted from bind irregularly to all nucleic acids, DNA and RNA, interfering with their function.

Similarly, p53 accumulation following CD also occurs within a narrow concentration range from no effect to a much higher p53 accumulation than observed after any dose of DD agent. This spike in accumulation is followed by the loss of p53 accumulation with further increases in CBL0137 concentration, likely due to the inhibition of general transcription by CBL0137 at doses higher than 1.5–2 μM (unpublished data). In contrast, p53 accumulation is not proportional to the degree of DD and stays at the same level at concentration of bleomycin spanning three orders of magnitude. Based on these findings, we propose that p53 transcription and translation are rate-limiting in response to DD and rather low. Therefore, even when only a few DNA breaks activate several DD-sensitive kinases, these kinases phosphorylate almost all of the newly synthesized p53, and further increases in the number of DNA breaks and activated kinases do not result in additional p53 stabilization.

We do not know exactly how and why p53 protein levels increase in response to CD. We previously described FACT-CK2-mediated phosphorylation of p53 on serine 392 (5); however, this phosphorylation cannot explain p53 accumulation. It is also unclear why CK2 phosphorylates p53 after FACT binds to disassembled nucleosomes. Although we cannot exclude other p53 modifications in response to DD that we have not been able to detect so far, p53 is likely significantly less modified in response to CD than DD.

Because DD is considered a major cellular stress with multiple sensors and signaling pathways involved, we were surprised that the transcriptional response to DD almost exclusively depends on p53. Although multiple studies have analyzed the transcriptional response to DD, those studies were performed using bulk RNA-seq that assessed the average response of millions of cells. At the single-cell level, where hundreds to thousands of cells are processed as individual replicates, we found that only the activation of very few classical p53 targets was statistically significant. DD inhibited many more genes in a p53-dependent manner; however, all the inhibited genes were expressed in cycling cells, with ‘E2F targets’, and ‘G2/M checkpoint,’ being the main enriched categories. These findings suggest that DD causes p53-dependent growth arrest by inducing *CDKN1A*, and the difference in gene expression is a difference between cycling and arrested cells. In support of this concept, the transcriptome of untreated, non-dividing normal fibroblasts is not different from that of p53-positive bleomycin-treated cells. However, we cannot exclude the role of p53 in suppressing the expression of some of these genes. There are still debates about the role of p53 in suppressing gene expression (42).

Transcriptional changes in response to CD differ from p53 signaling in response to DD. First, more genes are activated by CD than DD, and these genes are similarly activated in p53 wild-type and null cells. Moreover, a highly p53-specific reporter was activated by CD in the absence of p53. Thus, CD may act directly on the nucleosomes around the TSS of these genes, facilitating acts of transcription due to the presence of paused RNA polymerase II and a chromatin state characteristic of active transcription.

In p53 wild-type cells, we are most likely observing the combined effects of CD directly on nucleosomes or paused RNA polymerase II around the TSS of some genes and the accumulation of high levels of unmodified or weakly modified p53. Experiments using lentiviral delivery of p53 or inducible p53 expression demonstrated that poorly modified p53 is transcriptionally active (22,35). However, the input of p53 protein in the transcriptional response to CD does not seem essential because the degree of activation of endogenous p53 targets was not much different between p53-positive and p53-negative cells based on nascent RNA-seq data.

In summary, we identified several consequences of CD and compared them with those of DD. Although both DD and CD are cytotoxic, there are important differences in their effects on cells, such as the inability of CD to induce senescence. Moreover, both DD and CD are potent activators of p53 and p53-dependent transcription; however, only the response to DD is highly p53-dependent. The CD response is largely p53 independent. Based on our findings, we propose the following model: CD directly affects the promoters of stress-responsive genes, including p53 targets. Genes activated by CD have open chromatin and paused RNA polymerase II under basal conditions. Treatment with CD agents triggers the transcription of these genes even in the absence of specific transcription factors, such as p53.

## Data Availability

The following datasets are submitted to NCBI GEO datasets: scRNA-seq of NDF and HT1080 cells, untreated or treated with CBL0137 or bleomycin (accession number is pending), nascent RNA-seq of HT1080 cells untreated or treated with CBL0137 or bleomycin (accession number is pending), bulk RNA-seq data of untreated NDF cells (accession number is pending), MNase-seq data of HT1080 cells, untreated or treated with CBL0137 (accession number is pending).

## Acknowledgements

We are grateful for the help and support of the administrative office of the Cell Stress Biology Department of Roswell Park Comprehensive Cancer Center, specifically Mary Morgan, Sarah Tyrpak, Sarah Marcy and Bruce Specht. We thank the members of the shared resources facilities for help and advice with the design and processing of the experiments, specifically Dr. Prashant Singh, Director of the Genomics Shared Resource. We also thank Shawn Matott, a senior systems analyst, for the help with the High-Power Computer resources at RPCCC and University of Buffalo. Special thanks to Dr. Andrei Gudkov for the discussion of the manuscript and Dr. Catherine Burkhart and Burkhart Document Solutions for critical reading and scientific editing of the manuscript.

