## Supplementary Figures for "Comparison of cell response to chromatin and DNA damage"

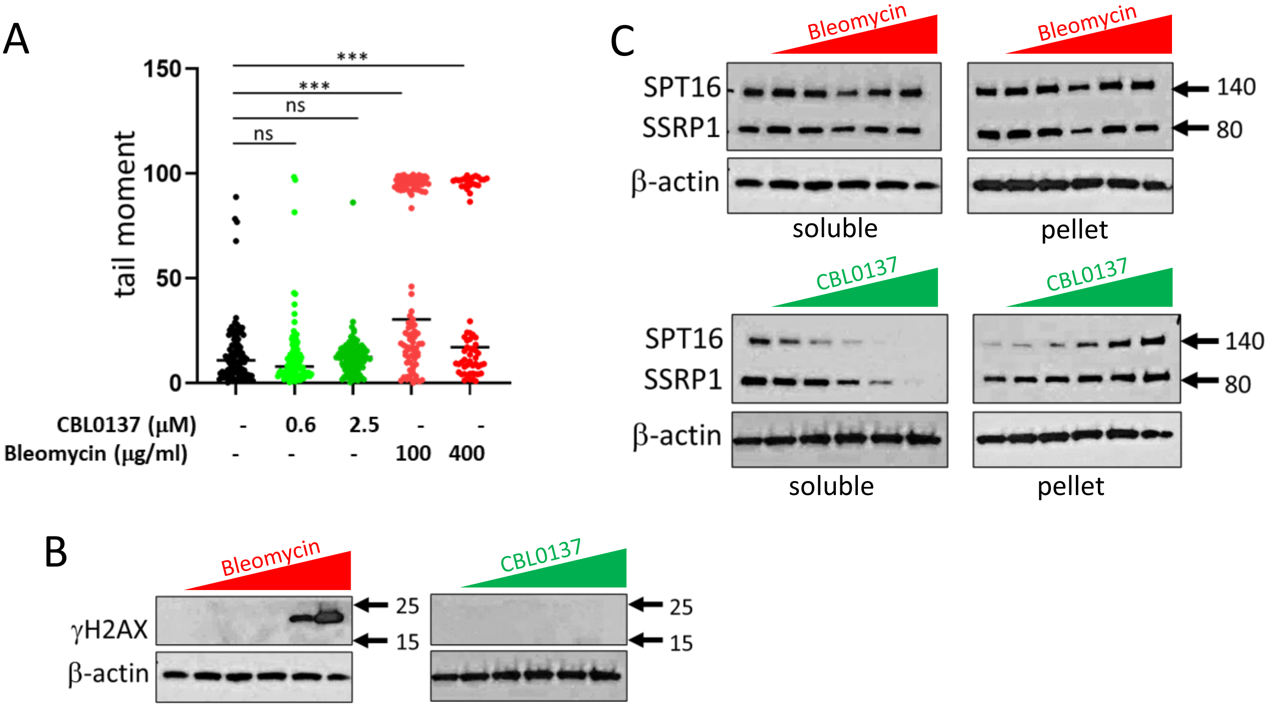


**Supplementary Figure S1. Comparison of DD and CD effects of bleomycin and CBL0137**. A. Alkali comet assay for cells treated with CBL0137 or bleomycin for 24 hours. Tail moment was assessed using the OpenComet plugin of ImageJ. A total of 100 randomly selected cells were evaluated. ***$p < 0.005$ by ANOVA. B. Western blotting of lysates of HT1080 cells treated with Bleomycin or CBL0137 for 24 hours and stained with γH2AX or β-actin antibody. C. Western blotting of soluble and pelleted protein fractions from HT1080 cells stained with antibodies against the indicated proteins. The soluble fraction represents the nucleoplasm, and the pelleted fraction represents chromatin. At B and C bleomycin was used at 63, 125, 250, 500, and 1000 µg/ml, and CBL0137 was evaluated at 0.325, 0.63, 1.25, 2.5, and 5 µM. The numbers to the right of the western blots are the positions of the protein size markers in kDa.

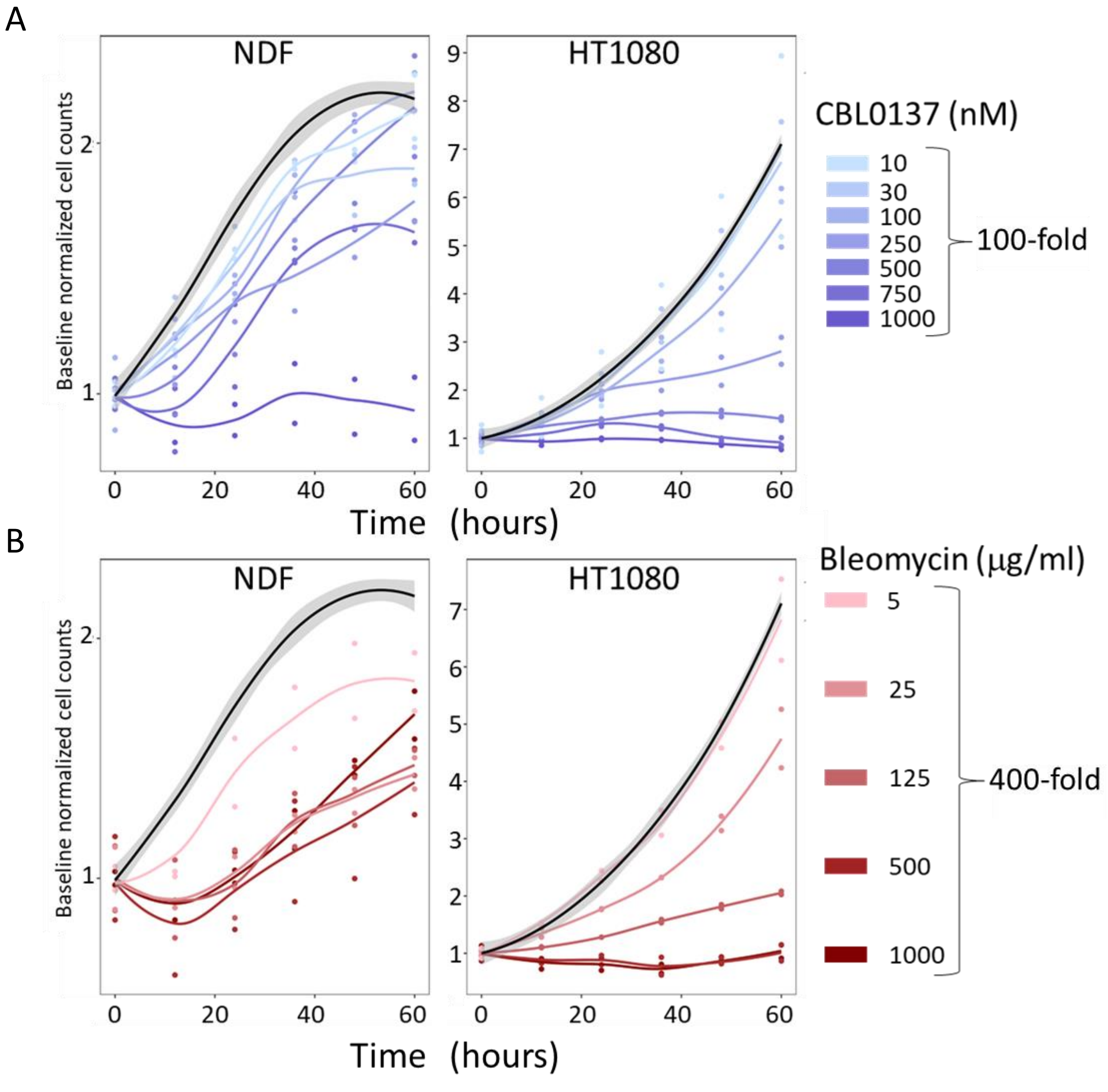


**Supplementary Figure S2. Time-dependent effects of CBL0137 and bleomycin on the growth of NDFs and HT1080 cells**. Cells were plated in 384-well plates in triplicate. The next day, the cells were treated with different doses of CBL0137 (A) or bleomycin (B). Cell number was evaluated by automatically counting of cells every 12 hr by a Cytation 5 Cell Imaging Multimode Reader using Gen5 ImagePrime software. Three replicates per condition are shown as dots. Untreated control cells are shown as a black line with a grey area representing the standard error for ten replicates. Cell counts were normalized to a time point immediately after drug addition (approximately 5 min).

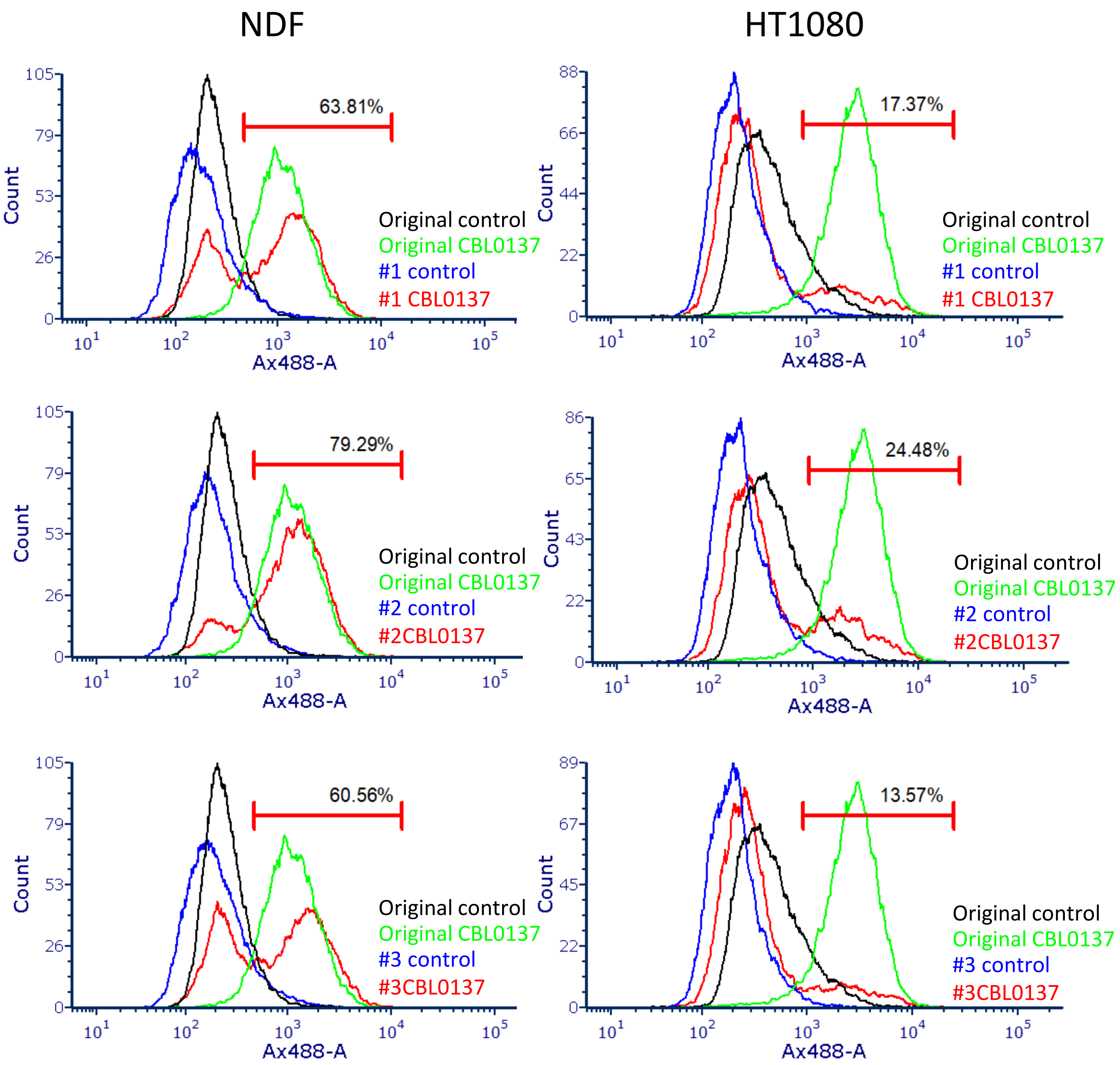


**Supplementary Figure S3. Assessment of the proportion of p53-negative cells following electroporation with Cas9 and gRNAs to *TP53* gene.** NDFs and HT1080 cells, parental and post-electroporation, were treated with 1 μM CBL0137 for 24 hr to induce p53. Treated and untreated cells were stained for p53 and analyzed by flow cytometry. Three slightly different electroporation regimens were used (#1, #2, and #3). The marker shows the position of the p53-positive cells in the parental cells after CBL0137 treatment. The percentage above each marker indicates the proportion of p53-positive cells after electroporation.

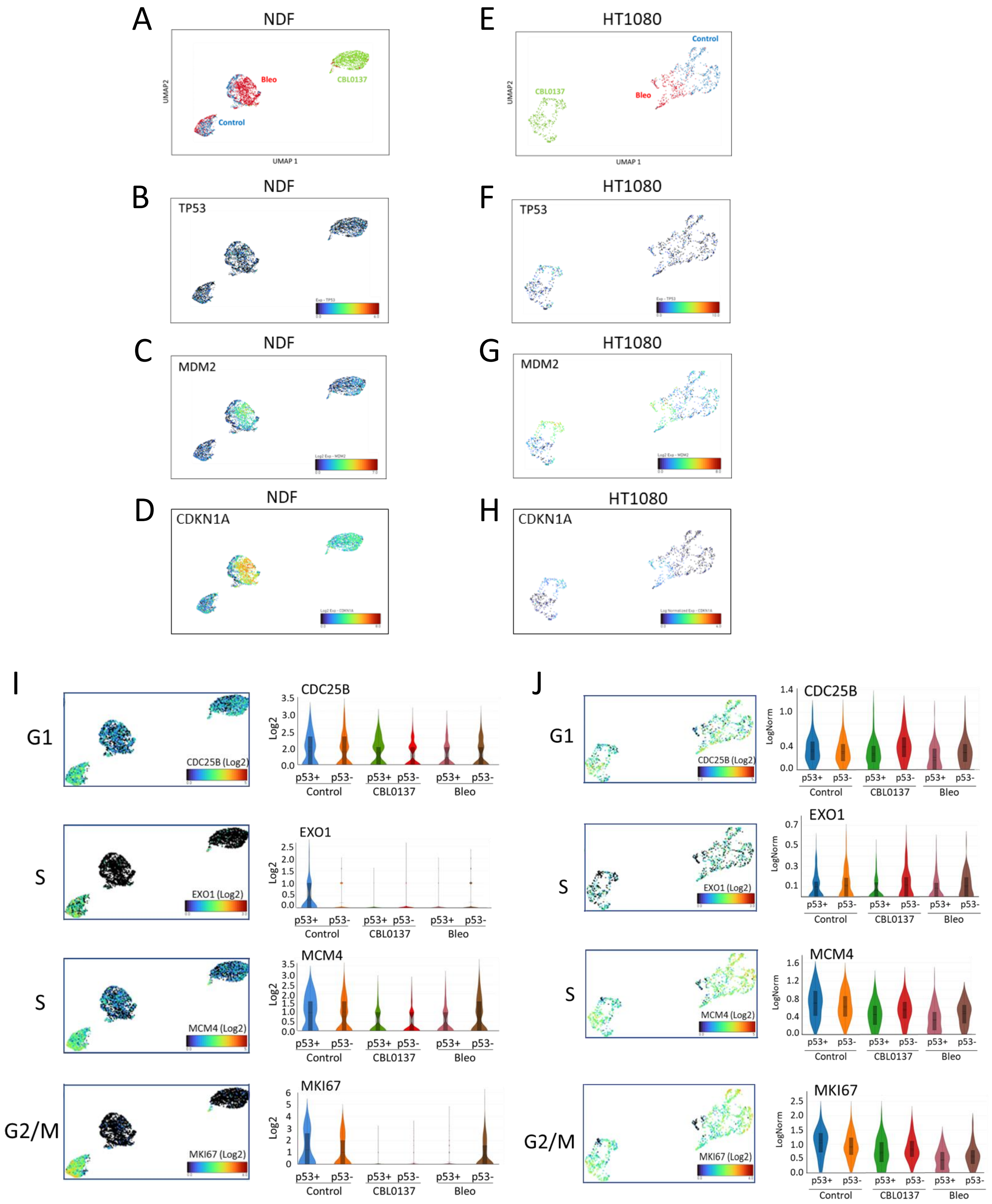


**Supplementary Figure S4. Analyses of scRNA-seq data from NDFs (A-D, I) and HT1080 (E-H, J) cells.** A-H. UMAP plots showing position of cells depending on treatment (A, E), or expression of the following genes, *TP53* (B, F), *MDM2* (C, G) and *CDKN1A* (D, H). I, J. UMAP and violin plots showing the expression of markers for the G1, S, and G2/M phases of the cell cycle in NDF (I) and NT1080 (J) cells under different conditions.

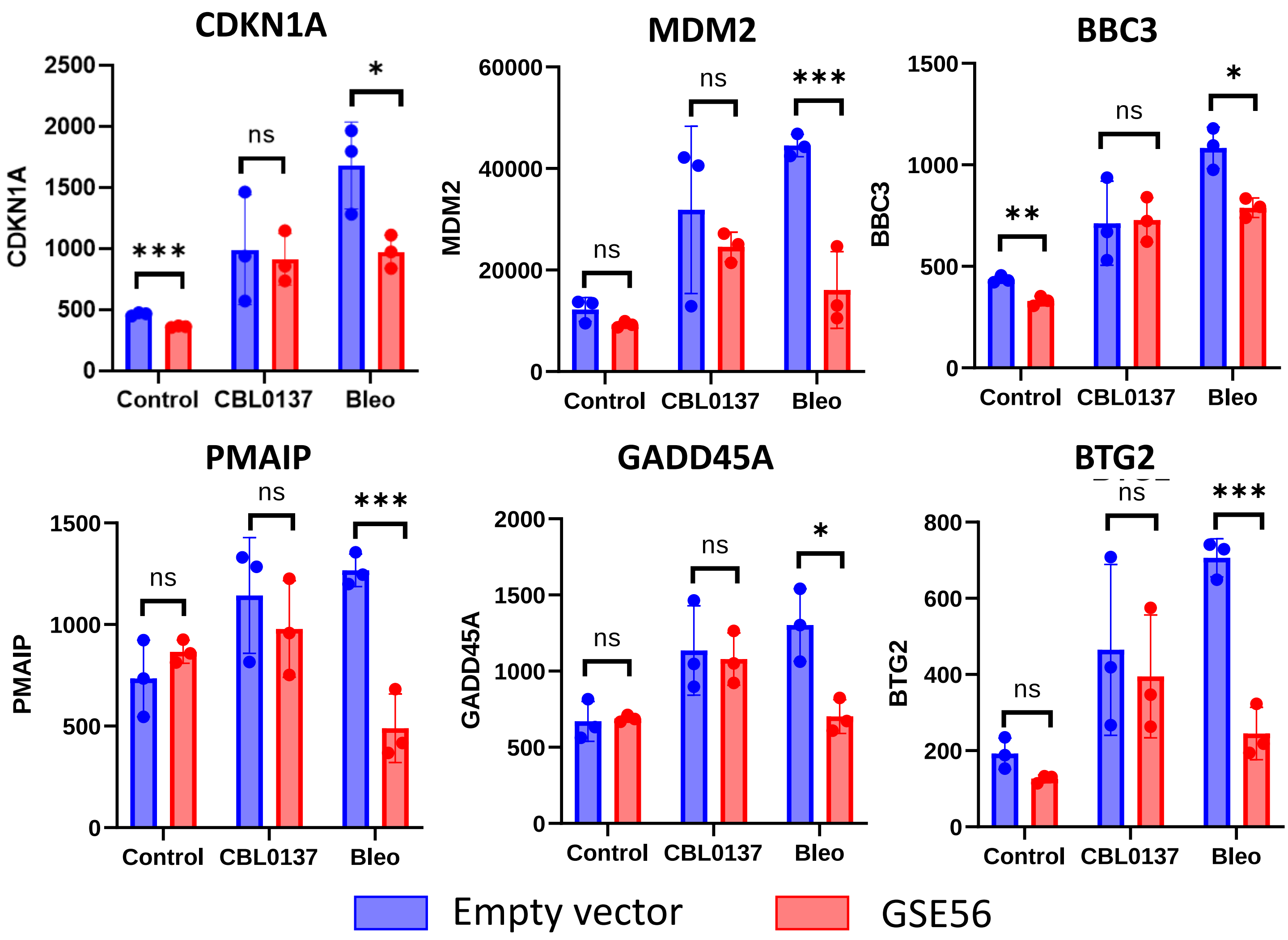


**Supplementary Figure S5. Transcription of p53 target genes in response to DD and CD.** Normalized data for nascent RNA-seq from HT1080 cells transduced with empty vector or p53 dominant-negative mutant GSE56 treated with 0.6 μM CBL0137 or 400 μg/ml bleomycin for 24 hr. Controls were left untreated. Data are presented as the mean ± SD (n = 3 replicates). *p < 0.05, **p < 0.01, ***p < 0.005, ns, not significant by the paired Student's t-test, functional p53 vs. inactive p53.

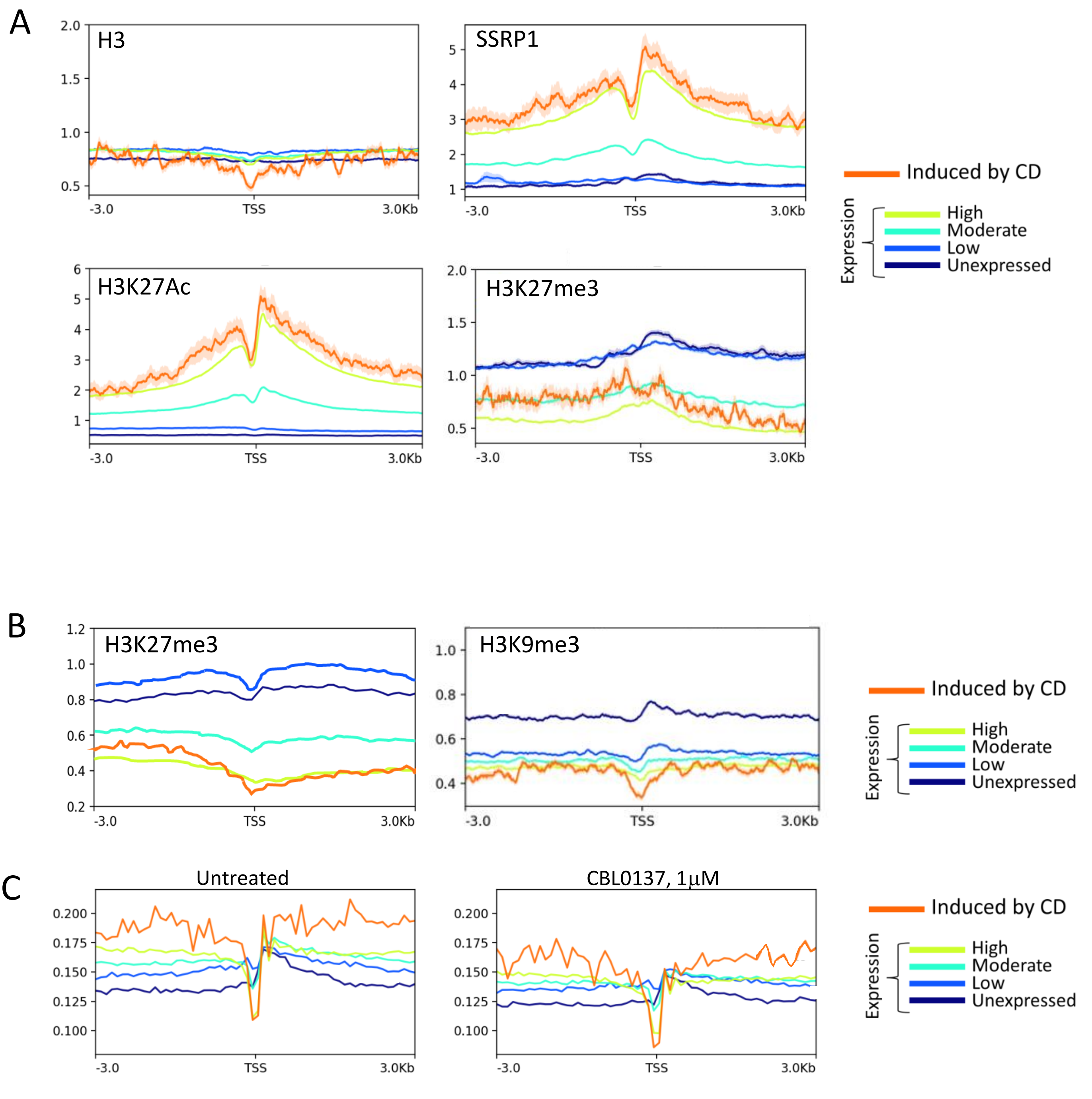


**Supplementary Figure S6. Comparison of chromatin state around TSS of genes induced by CD and genes transcribed at different levels. A.** Metagene profiles of total histone H3 (as a proxy of nucleosome occupancy), SSRP1 subunit of FACT complex (as a proxy of active transcription), and two posttranslational modifications of histone H3, H3K27 acetylation (marker of active transcription) and H3K27 trimethylation (marker of repressed transcription) in untreated HT1080 cells. B. Metagene profiles of repressive histone post-translational modifications in untreated NDF cells . B. Metagene profiles of DNA protected from MNase digestion in untreated HT1080 cells and HT1080 cells treated with CBL0137 for 1 hour.
